# Structure and regulation of full-length human leucine-rich repeat kinase 1

**DOI:** 10.1101/2022.12.21.521433

**Authors:** R.D. Metcalfe, J.A. Martinez Fiesco, P. Zhang

## Abstract

The two human leucine-rich repeat kinases (LRRKs), LRRK1 and LRRK2 are large and unusually complex multi-domain kinases which serve to regulate fundamental cellular processes, with both implicated in human disease. Despite recent experimentally determined structures of LRRK2, the structure and exact molecular mechanisms regulating the activity of the LRRK1, and differences in the regulation of LRRK1 and LRRK2 remain unclear. Here, we report a cryo-EM structure of the LRRK1 monomer, and a lower-resolution cryo-EM map of the LRRK1 dimer. The monomer structure, in which the kinase is in an inactive conformation, reveals key interdomain interfaces which serve to control kinase activity, which are further validated experimentally. Both the LRRK1 monomer and dimer are structurally distinct compared to LRRK2, implying functional differences in both proteins. Overall, our results provide new structural insights into the activation of the human LRRKs for understanding the physiology and pathology of these proteins.

## Introduction

Leucine-rich repeat kinase (LRRK) 1 is a multidomain protein which, near-uniquely, contains both a Ras-like GTPase domain and kinase domain in the same polypeptide chain. The two human LRRKs, LRRK1 and LRRK2 was first identified in 2002 in a genome screen for unidentified human kinases^1^. Shortly after, mutations in LRRK2 were shown to be pathogenetic, causing both familial and sporadic Parkinson’s disease (PD)^2,3^, which has led to intense investigation of the biological role of LRRK2, and the development of LRRK2-specific kinase inhibitors^4,5^. Despite this, the exact physiological roles of both LRRK1 and LRRK2 is still unclear. Both LRRK1 and LRRK2 phosphorylate distinct subsets of Rab GTPases on a conserved serine or threonine, which results in the alteration of interactions between the Rab proteins and their effectors^6^.

The biological roles of LRRK1 are less well-established compared to LRRK2. LRRK1 and LRRK2 interact with distinct proteins and have different substrates and are thus functionally non-redundant proteins^7,8^. LRRK1 does not phosphorylate the LRRK2 substrates Rab10 and Rab8, instead phosphorylating Rab7, a key player in lysosomal biogenesis and trafficking^7,9^. Related to this, LRRK1 has roles in regulating mitophagy and autophagy^10,11^. Likewise, LRRK1 has a role in regulating the trafficking and lysosomal degradation of the epidermal growth factor receptor (EGFR)^12,13^. LRRK1 is itself tyrosine-phosphorylated by the EGFR, which places LRRK1 in a pathway regulating EGFR trafficking^13^. LRRK1 has a well-established role in bone development^14^, with LRRK1 knockout mice exhibiting severe osteoporosis^15^. A genetic disease, osteosclerotic metaphyseal dysplasia (OSMD), which is characterized by severe bone abnormalities and osteoporosis, is caused by mutations in LRRK1 which eliminate LRRK1 kinase activity^7,16–18^. The Rho-family small GTPase Rac1, which has a role in regulating the actin cytoskeleton in osteoclasts^19^, is phosphorylated by LRRK1^20^, suggesting a mechanism for the link between the loss of LRRK1 activity and OSMD.

Both LRRK1 and LRRK2 are exceptionally large proteins (230-280 kDa, 2000-2500 residues), with complex domain structures^21,22^. LRRK1 consists of N-terminal ankyrin repeats, leucine-rich repeats, and C-terminal Roc (Ras-of-complex) GTPase domain, COR (C-terminal of Roc) scaffolding domain, kinase domain and a WD40 repeat domain at the C-terminus, with LRRK2 additionally having armadillo repeats N-terminal of the ankyrin repeats. The Roc-COR tandem domain is characteristic of the Roco family of proteins, which is conserved in all domains of life^23^. The LRRKs are a subset of Roco proteins, which are limited to animals^24^. Early biochemical analysis showed that both LRRKs are functional kinases, however biochemical characterization, and structural analysis was hindered by the size, flexibility, and low recombinant expression levels of both proteins^22,25–29^. Recently, cryo-electron microscopy (cryo-EM) structures have been reported of the catalytic C-terminal half of LRRK2^30^, full-length inactive LRRK2^31^, the catalytic half of LRRK2 engaged with microtubules^32^ and the active LRRK2 bound to a Rab protein^33^. A low-resolution cryo-EM map of the catalytic half of LRRK1 suggests that the arrangement of the catalytic domains is conserved between LRRK1 and LRRK2^32^, although a detailed structural analysis of the full-length LRRK1 has so far not been possible.

Here, we report the cryo-EM structure of the monomeric full-length LRRK1. Our structure explains several regions known to regulate LRRK1 kinase activity. The structure reveals notable structural differences between LRRK1 and LRRK2, most significantly in the position and structural dynamics in the leucine-rich repeats, but additionally in interdomain contacts between the kinase and Roc domains. Our results provide a structural framework for examining LRRK1 biology and allow the future dissection of unique and universal mechanisms of the activation and regulation of both LRRK1 and LRRK2.

## Results and Discussion

### Purification and initial characterization of LRRK1

We purified full-length human LRRK1 (residues 1-2015) from baculovirus-infected insect cells, using sequential Flag-affinity and gel filtration chromatography (Supplementary Figure 1A-B). The initial affinity purified protein eluted from the gel filtration column as a complex mixture of monomeric, dimeric, and larger species (Supplementary Figure 1A-B). We determined the solution mass of the putative ‘monomer’ and ‘dimer’ fractions using mass photometry^34^ (MP) (Figure 1A-B), identifying species with masses consistent with the presence of the LRRK1 monomer (229 kDa) and dimer (458 kDa) in both fractions (Figure 1A-B), with the monomer fraction predominately containing monomers, and the dimer fraction containing both monomers and dimers at approximately a 1:2 ratio, alongside other higher-order species. Multi-angle light scattering (MALS) data collected on the monomer fraction shows that the mass is 248 kDa, again consistent with the expected mass of the LRRK1 monomer (Supplementary Figure 1C). These results show that LRRK1 (1-2015) exists as both a monomer and dimer in solution and does not preferentially form either species.

**Figure 1:**
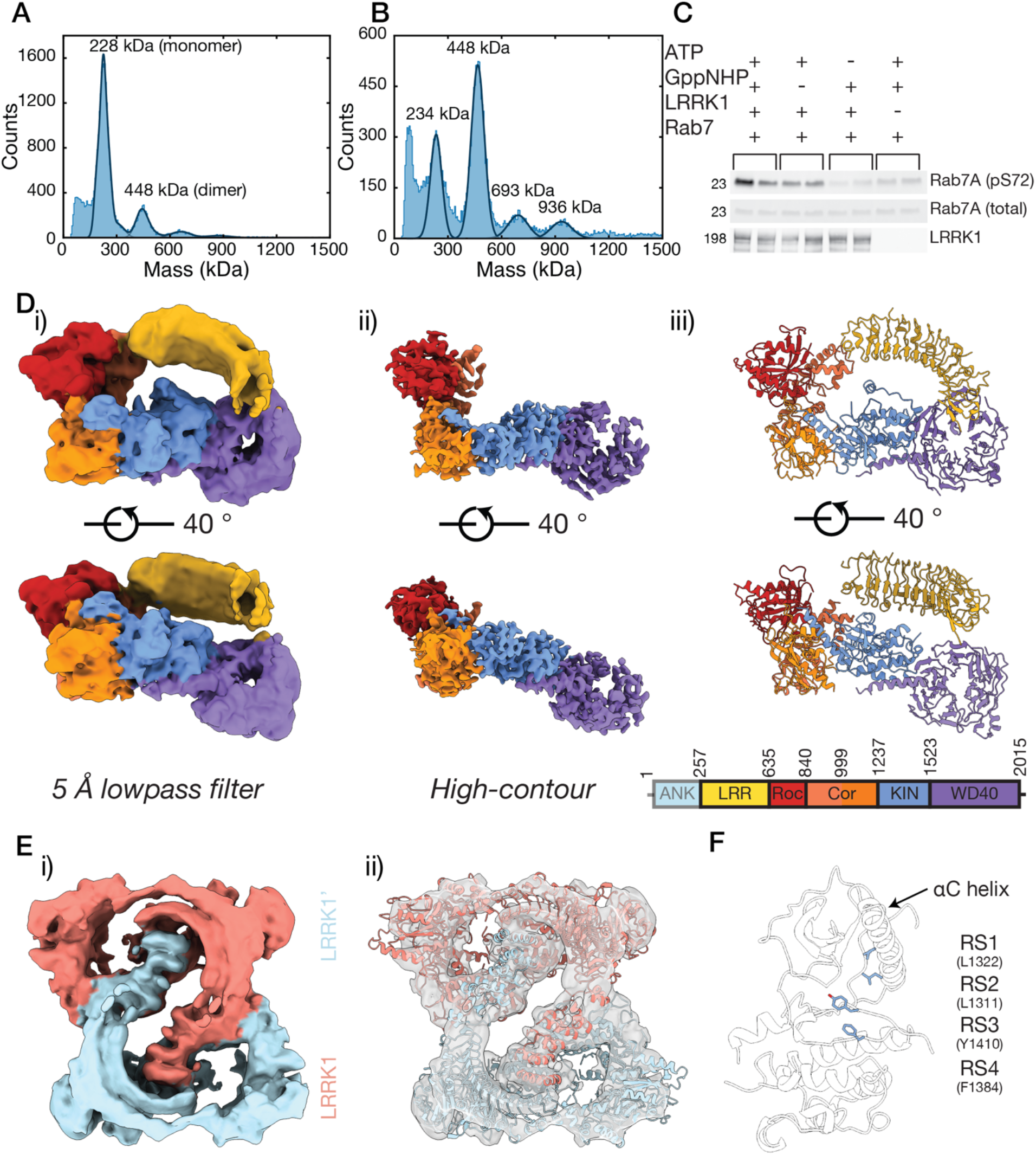
Structure of LRRK1 and the inactive kinase domain of LRRK1. A) Mass photometry mass distributions for the LRRK1 monomer, and B) the LRRK1 dimer fractions. C) Western blot showing phosphorylation of the LRRK1 substrate Rab7A by purified LRRK1 (see Supplementary Figure 1 for complete membrane images). D) Cryo-EM map and model of the LRRK1 monomer, i) map low-pass filtered to 5 Å, ii) map rendered at a high-contour level, showing detail in the C-terminal domains, iii) atomic model of the LRRK1 monomer, colored according to the schematic. E) Cryo-EM analysis of the LRRK1 dimer, i) cryo-EM map of the LRRK1 dimer, ii) two copies of the LRRK1 monomer fit in the LRRK1 dimer map. F) The broken kinase R-Spine (RS) in the kinase domain in the LRRK1 monomer, indicating that the kinase domain is in the inactive conformation.

We observed robust Rab7A phosphorylation by the purified LRRK1 monomer using a Western blot assay (Figure 1C, Supplementary Figure 1D), confirming that the purified LRRK1 can phosphorylate its substrate.

### Structure of full-length human LRRK1

We subsequently undertook single particle cryo-EM of LRRK1 to determine the structure (Figure 1D-E, Supplementary Figure 2, see Methods). We collected data on grids prepared from both LRRK1 monomer and LRRK1 dimer fractions and processed the data together, as the LRRK1 dimer datasets contained a substantial number of LRRK1 monomer particles, as would be anticipated from the MP measurements (Figure 1B, Supplementary Figure 2A-B). 2D classification revealed several classes with the domains of LRRK1 clearly visible, and several classes with clear twofold symmetry, corresponding to the LRRK1 dimer (Supplementary Figure 2A-B). Subsequent processing resulted in a 3.9 Å resolution map of the LRRK1 monomer and a 6.4 Å map of the LRRK1 dimer (Figure 1D-E, Supplementary Figure 2C, Supplementary Figure 3A-C, Supplementary Figure 4A-B). Local refinement improved the resolution of the C-terminal Roc-COR/kinase/WD40 catalytic domains to 3.8 Å in the LRRK1 monomer map (Supplementary Figure 2C, Supplementary Figure 3B-C). In the monomer map, large sidechains are visible and secondary-structure elements are well defined, α-helices clearly visible, and β-strands separated, consistent with a map reconstructed at this resolution (Supplementary Figure 3D). The definition within the LRRs is significantly poorer (Figure 1Di-ii, Supplementary Figure 3A-B).

The overall structure of the LRRK1 monomer is ‘O-shaped’. The structure is relatively compact, with the Roc, COR, kinase and WD40 domains forming an ‘J-shape’. The leucine-rich repeats bridge the Roc and WD40 domains (Figure 1D), although they do not form direct contacts with the WD40 domain. The ankyrin repeats are not visible in the density, implying that they are flexible in the monomer. We purified LRRK1 in the absence of any nucleotide and did not add any nucleotide prior to cryo-EM grid preparation. We did not observe any density for an adenosine nucleotide in the kinase active site. We did include GDP in the atomic model, as we did observe density for GDP in the Roc active site, likely endogenous GDP (Supplementary Figure 5A-B). Reinforcing this observation, the Switch-I GTPase motif, which responds to guanosine nucleotide binding^35^, is unambiguously in a conformation consistent with GDP binding (Supplementary Figure 5A-B).

Two copies of the LRRK1 monomer fit in the LRRK1 dimer map (Figure 1Eii). Individual domains of the LRRK1 monomer are visible within the density (Figure 1Eii). Additional helical density is visible above the kinase domain in the map, adjacent to the LRRs, which we assigned to the ankyrin repeats (Supplementary Figure 4C). The ankyrin repeats form the dimerization interface between the two LRRK1 monomers. The LRRs and ankyrin repeats block the kinase active site of the adjacent LRRK1 molecule in the dimer, implying that the dimer is intrinsically inactive (Figure 1E, Supplementary Figure 4A), however the low resolution of the map prohibits a detailed analysis of the dimerization interface (Figure 1E, Supplementary Figure 4).

In the structure, the LRRK1 kinase domain is in an inactive conformation, and packs closely to the COR-B domain (Figure 1B,F). The kinase regulatory (R)-spine^36^ is broken, the αC helix is in the ‘out’ position, and the conserved K1270/E1307 ion pair is broken (between the αC helix and N-lobe, equivalent to the K72/E91 ion pair in PKA), all markers of an inactive kinase (Figure 1F). The kinase activation loop is well-defined in the density (Supplementary Figure 3D), with the conserved DFG (DYG in LRRKs) residue D1409 in an ‘out’ conformation, pointed away from the ATP binding site.

### Comparison of the structures of LRRK1 and LRRK2 reveal striking difference in position of LRRs

Recent structures of LRRK2 allow the comparison of experimental structures of LRRK1 and LRRK2^30–33^ (Figure 2A). The most striking difference between the two proteins is in the position of the LRRs (Figure 2A). In LRRK2, the LRRs and ANK repeats cover and occlude the kinase domain, presumably serving to regulate substrate access (Figure 2B) with the additional N-terminal domains in LRRK2 projecting beyond the kinase-LRR ‘core’ of the protein, with the ‘hinge helix’ in the LRRs forming an interface between the ankyrin, LRR and WD40 domains^31^. The overall arraignment of the C-terminal catalytic domains is conserved between the two proteins, and the overall structure of the kinase domain in LRRK1 and both the determined inactive LRRK2 kinase structures are very similar, with the major structural difference being an extended αC helix in LRRK1 (Figure 2C). Relative to the kinase, the Roc domain is displaced by ~10 Å between the two proteins, which accommodates sterically the more compact position of the LRRs in LRRK2 (Figure 2D). The curvature of the LRRs is very similar between the two proteins. The position of the LRRs in LRRK2 implies the requirement for rearrangement of the LRRs and N-terminal domains to permit substrate access to the kinase in LRRK2. In contrast, in LRRK1 the LRRs do not form interfaces with the remainder of the protein and do not occlude the kinase domain (Figure 1).

**Figure 2:**
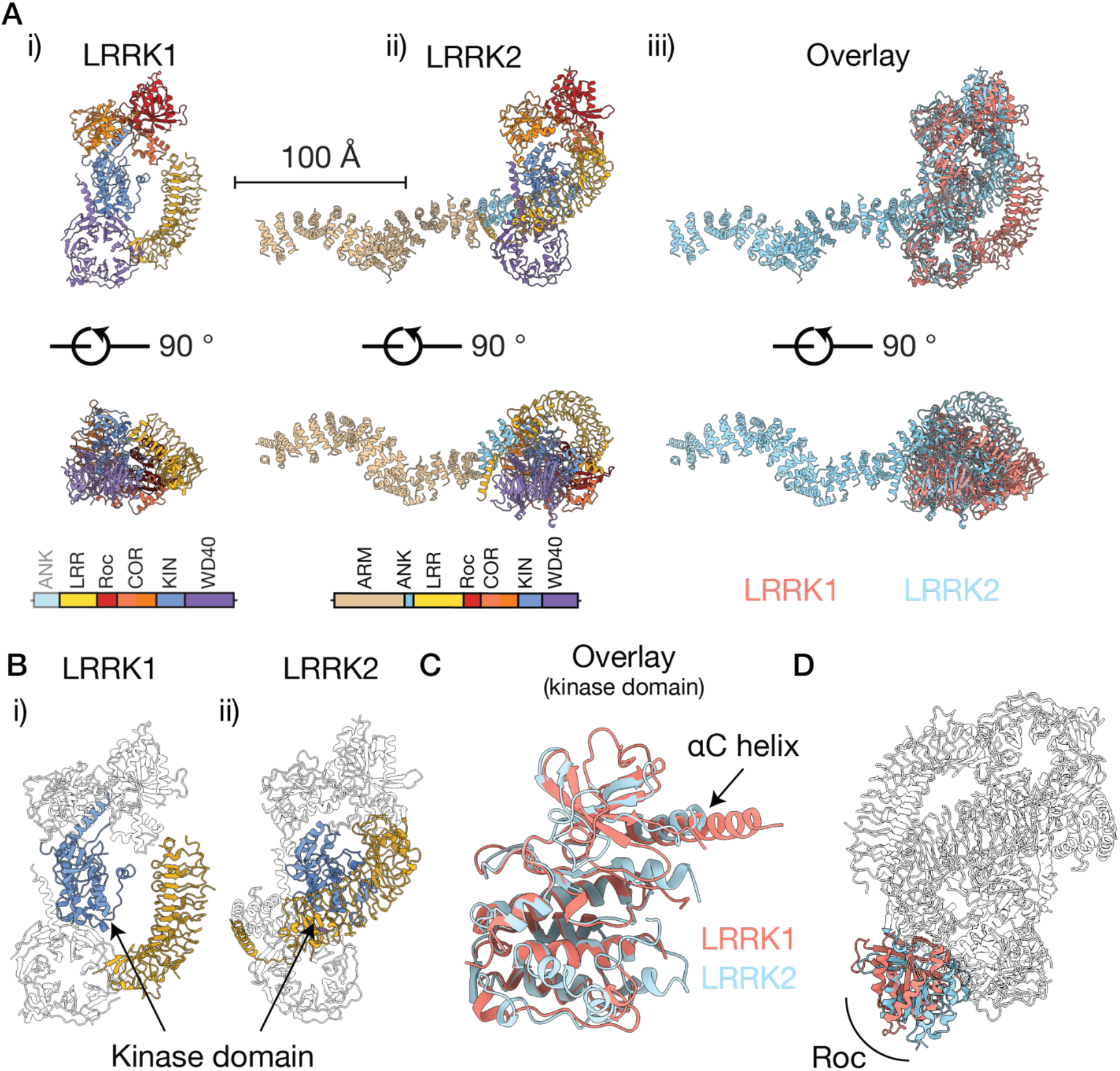
Structural comparison of the LRRK1 and LRRK2 monomers. A) The structure of the LRRK1 monomer, i) and the LRRK2 monomer^31^ (PDB: 7LHW, EMD-23352, residues 1-543 from predicted Alphafold^41,54^ model AF-Q5S007 ii), and an overlay, iii). Structures are displayed on the same scale. B) Relative position of the LRRs and the kinase domain for LRRK1, i) and LRRK2, ii), showing that the kinase domain in LRRK2 is occluded by the LRRs. C) Overlay of the kinase domain of LRRK1 and LRRK2. D) Structures aligned on the kinase domain, highlighting the shifted position of the Roc domain between the two proteins, which accommodates the altered position of the LRRs.

The kinase domain is occluded in both the LRRK1 and LRRK2 dimers (Supplementary Figure 4D-E). In the LRRK1 dimer, the kinase domain is occluded by the ankyrin repeats from the neighboring LRRK1 molecule in the dimer, in LRRK2, the kinase domain is occluded by the LRRs of the same monomer. This shows that the current LRRK dimer structures represent intrinsically inactive forms of the molecule.

### Structural scaffolds in the kinase and GTPase Roc-COR domains control LRRK1 activity

Two structural scaffolds in the COR-B and kinase domains of LRRK1 serve to link the two domains and tightly regulate the LRRK1 kinase domain through controlling the kinase inactive-to-active transition. First, in LRRK1, the unusually long αC helix, which is approximately twice as long compared to the αC helix in LRRK2 and other comparable kinases, acting as a structural scaffold linking the kinase N-lobe and the COR-B domain (Figure 3A-C, Supplementary Figure 6). In LRRK1, the kinase αC helix sits between the kinase N-lobe and the COR domain, with the extended helix forming several contacts with the COR-B domain that are unique to LRRK1 (Figure 3B-C). Typically, the αC helix transitions from an ‘out’ to ‘in’ position on kinase activation, allowing an overall rearrangement of the kinase core to allow substrate phosphorylation^37^. Interactions involving the αC helix often serve to be a key regulator of kinase activity in multidomain kinases or kinase complexes^38,39^.

**Figure 3:**
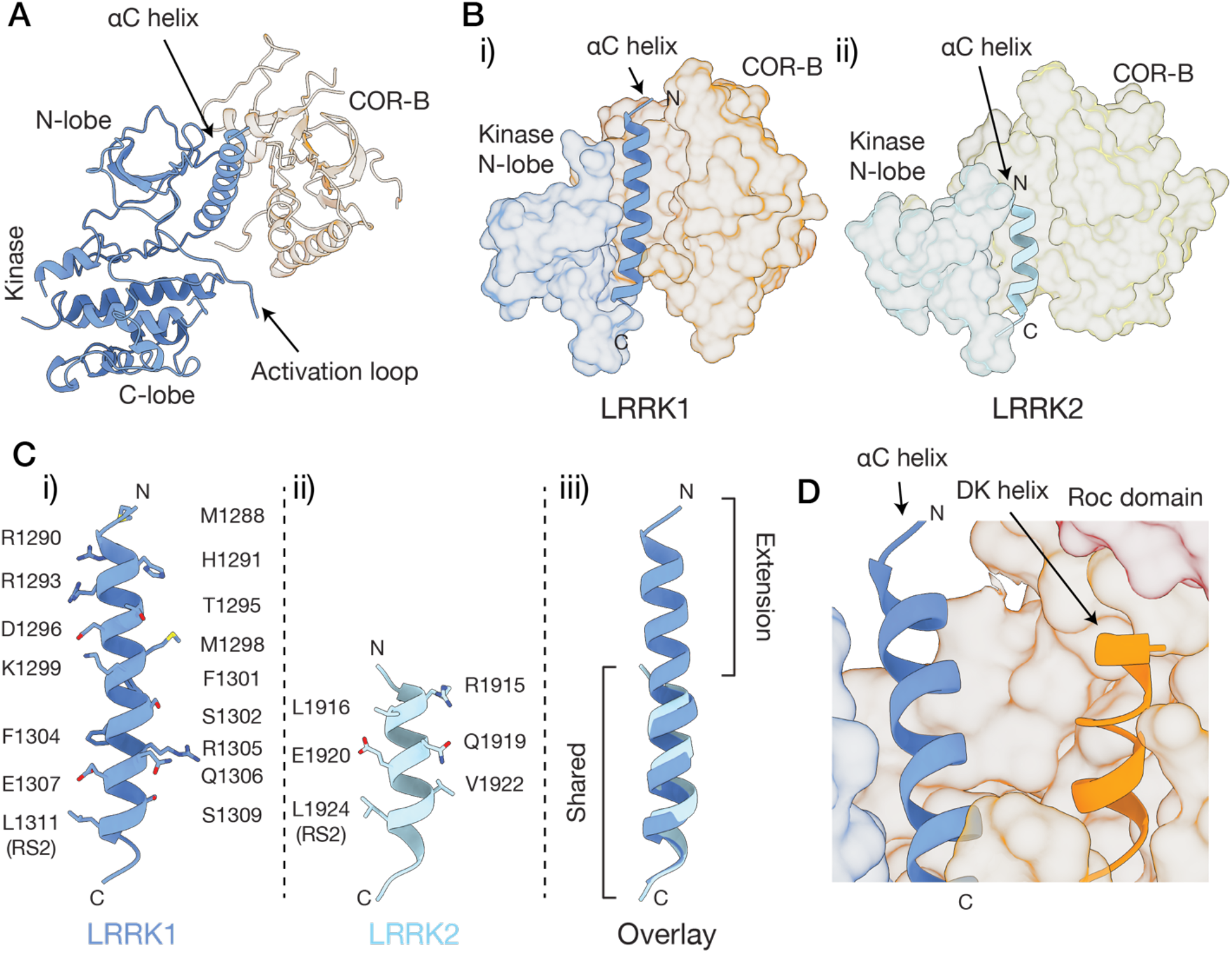
Interdomain interfaces regulating the LRRK1 kinase domain. A) The interface formed by the LRRK1 kinase and the COR-B domain. B) The interface formed between the kinase N-lobe, αC helix and the COR-B domain in LRRK1, i) and LRRK2 ii). C) The αC helix in LRRK1, i), LRRK2 ii) and an overlay, iii), with residues along the LRRK1 and LRRK2 αC helix indicated in i) and ii), the shared region, and the unique LRRK1 extension indicated in iii) (see also Supplementary Figure 6). D) The position of the COR-B DK helix relative to the kinase αC helix and the Roc domain (shaded red).

The second key scaffold In the LRRK1 catalytic domains is the DK helix (also known as the COR-B helix, residues 1132-1144). The DK helix faces and sitting at approximately at 30° angle to the αC helix. In contrast to the inactive LRRK2 structure, where the DK helix forms several electrostatic contacts with the αC helix^31,40^, the LRRK1 DK helix is shifted by ~3 Å relative to LRRK2 and thus does not form extensive electrostatic contacts with the αC helix (Figure 3D). Hydrophobic residues on the face of the DK helix opposite the αC helix pack tightly with COR-B domain, stabilizing this position of the helix (Figure 3D). The C-terminal end of the DK helix sits against the Roc domain, creating an interface which has been shown to be key for LRRK2 regulation^40^.

We tested the effect of a set of mutations at the αC helix/COR-B interface and the Roc/COR-B interfaces using an *in vitro* kinase assay, measuring Rab7A phosphorylation by purified, recombinant LRRK1 as a readout (see Methods). We expressed a set of mutations that straddle both the region of the αC helix, which is conserved and the extension which is unique to LRRK1 (Figure 3B-C, Figure 4A, Supplementary Figure 6). First, we studied the R1305A and F1301A mutations, which form part of the conserved αC helix/COR-B interface (Figure 3C, Figure 4A-B). Both mutants express well *in vitro.* The F1301A does not alter Rab7A phosphorylation (Figure 4A, p=0.3311 relative to WT, see Supplementary Figure 7A-C throughout for complete membrane images). The R1305A mutation decreased Rab7A phosphorylation to the same level as the K1270M kinase-dead mutant (Figure 4A, p=0.0018 relative to WT, p=0.8129 relative to K1270M). R1305A is thus clearly involved in LRRK1 activation. In the inactive kinase state, R1305 protrudes into a small pocket formed by the COR-B domain, stabilizing it. Although an experimentally-determined active-kinase structure of LRRK1 is not available, the predicted Alphafold^41^ model of LRRK1 has the kinase in an active conformation, and is reminiscent of the LRRK2 active conformation^33^. In the Alphafold model, the COR-B DK helix undergoes a large rearrangement, resulting in the formation of a hydrogen bond between D1135 in the DK helix and R1305 (Figure 4B-C), a residue which is ~6 Å from the αC helix in the inactive conformation, suggesting R1305 has a key role for controlling the LRKR1 inactive-to-active transition.

**Figure 4:**
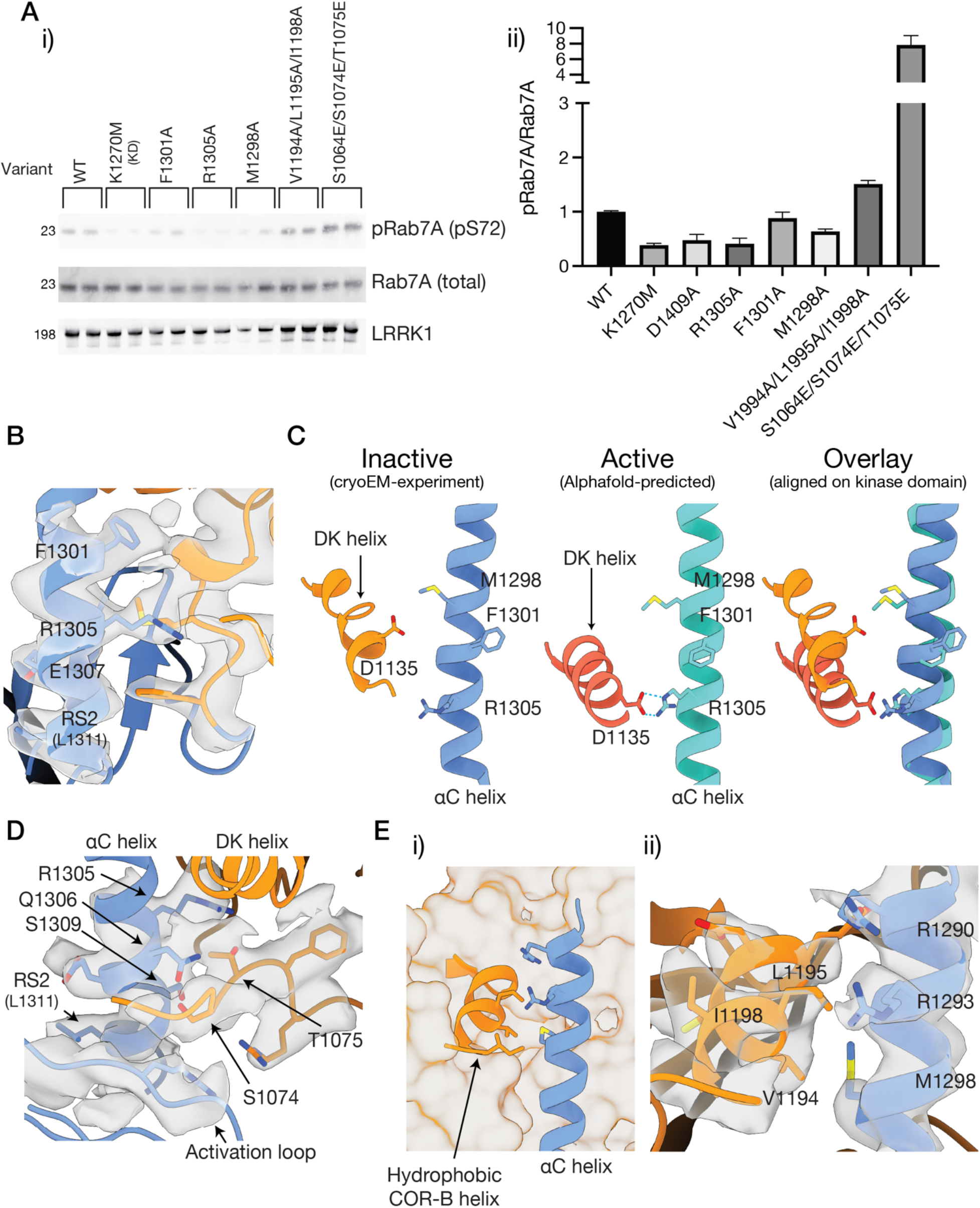
The kinase αC helix/COR-B interface. A) Effect of mutations in the kinase αC helix/COR-B interface on Rab7A phosphorylation by recombinant LRRK1, i) Western blot showing Rab7 phosphorylation, ii) quantification of multiple kinase assays from at least three independent protein preparations (see Methods, Supplementary Figure 7A for complete membrane images, values are mean ± SEM). B) Cryo-EM density supporting the position of F1301 and R1305 at the conserved, C-terminal end of the αC helix. C) Structural changes between the cryo-EM model, which is in the inactive state, and the active-state Alphafold model, highlighting a predicted interaction between R1305 and D1135 in the COR-B DK helix. D) Cryo-EM density supporting the position of S1074 and T1075 at the base of the αC helix. E) Interaction between the extended αC helix and a hydrophobic helix in the COR-B domain, i) showing the position of the helix, ii) cryo-EM density supporting this interface.

Close to the conserved αC helix region, a loop containing two residues (S1074, T1075) which are phosphorylated by PKC^42^ sit between the base of the DK helix and the αC helix, and close to the activation loop (Figure 4D). PKC phosphorylation of these residues results in the activation of the LRRK1 kinase^42^. The R-Spine residue L1311 sits on the opposite face of the helix facing the loop (Figure 4D), showing that the residues phosphorylated by PKC are at a key regulatory interface in the protein. In the unphosphorylated state, Q1306 and S1309 in the αC helix face the loop and interact with S1074 and T1075, stabilizing the ‘inactive’ position of this loop. This loop is poorly resolved in the density beyond N1071 which is the last residue included in the model, and the third PKC phosphorylation site (S1064) is thus not included in the model. The triple-phosphomimetic mutation, S1064E/S1074E/T1075E mimics PKC phosphorylation, *in cellulo* increasing LRRK1 Rab7A phosphorylation, albeit not to the levels by PKC phosphorylation^42^. Likewise, we observed that the recombinant triple-phosphomimetic increases Rab7A phosphorylation 5-fold over recombinant WT LRRK1 (Figure 4A, p=0.0024). Phosphorylation of these residues would be expected to result in a rearrangement of this critical regulatory interface involving the αC helix, the activation loop and the COR-B domain allowing kinase activation.

Next, we studied two mutations which form part of the ‘extended’ αC helix in LRRK1, M1298A and R1290A/R1293A (Figure 3Ci). Both mutants expressed poorly relative to the WT, K1270M, F1301A and R1305A mutants, we were unable to characterize the R1290A/R1293A doublemutant. The M1298A mutation modestly decreases LRRK1 Rab7A phosphorylation, but not to the level of the K1270M kinase-dead mutant (Figure 4A, p= 0.0002 relative to WT, 0.0009 relative to K1270M), however some caution should be taken in interpreting this result as the low expression level of M1298A made accurately measuring the kinase concentration difficult, regardless, M1298A does not abolish kinase activity. M1298 does not form any contacts with the COR-B domain in either the experimentally determined inactive state structure or the Alphafold model (Figure 4B-C). We additionally tested the effect of the removal of hydrophobic residues in a short α helix in the COR-B domain which contacts the extended αC helix (V1994A/L1195A/I1198A, Figure 4E). This mutation modestly but consistently increases Rab7A phosphorylation by ~1.5 fold (Figure 4A, p=0.0003 relative to WT). This shows that the unique kinase αC helix/COR-B contacts formed by the LRRK1 αC helix have a role in stabilizing the inactive state of the LRRK1 kinase. The fact the M1298A and R1290A/R1293A at the extended αC helix region expressed poorly suggest that they may play a role in protein stabilization in general, or in ensuring correct protein folding.

We next tested the effect a set of mutations in the Roc/COR-B interface formed by the DK helix (Figure 5A). There are known activating mutations in this interface in both LRRK1 and LRRK2^7,43,44^. In particular, mutation of K746, which sits at the Roc/COR-B interface (Figure 5B), is a known activating LRRK1 mutation^7^. In agreement with this, in our *in vitro* assay, the K746G mutation substaintially increased Rab7A phosphorlyation by 12-fold (Figure 5A, p=0.0148 relative to WT). In the cryo-EM model, K746 sits above the N-terminal end of the helix (Figure 5B-C). In the Alphafold active-state model, the DK helix undergoes a 15 ° shift relative to the inactive-state cryo-EM structure, resulting in the formation of extensive contacts between the DK helix and the αC helix, and placing the DK helix close to the kinase activation loop (Figure 5B-D), additionally placing K746 further away from the DK helix (Figure 5C). We tested the effect of several additional mutations in this interface, R1030A, W1144A, a W1144A/K746G doublemutant and R1034A. Except for the R1030A mutant, which did not express to levels sufficient to allow biochemical characterization, all mutations reduced or abolished Rab7A phosphorylation (Figure 5A). W1144 sits atop the DK helix, and, in the inactive state, is located close to K746 (Figure 5C). In the Alphafold model of the active state, the shift in the DK helix places W1144 close to the kinase activation loop, analogous to a conformational change seen in the active state of LRRK2^33^. The W1144A mutation reduces Rab7A phosphorylation to the same level as the K1270M kinase-dead mutant (Figure 5A, p=0.0004 relative to WT, 0.8089 relative to K1270M), as does the W1144A/K746G double-mutant (Figure 5A, p=0.0001 relative to WT, 0.0502 relative to K1270M). The R1034A mutation likewise reduces Rab7A phosphorylation, to the level of the K1270M kinase-dead mutant (Figure 5A, p=0.0001 relative to WT, p=0.1146 relative to K1270M). In the cryo-EM model of the inactive structure, R1034 sits alongside the DK helix, in both the cryo-EM model of the inactive state, and the Alphafold model of the active state, R1034 interacts with the DK helix, stabilizing the active conformation, rationalizing the kinase-inactivating properties of the R1034A mutant (Figure 5C). Thus, the Roc/COR-B interface through the DK helix is key for regulating LRRK1 kinase activity, despite its distant location from the key kinase regulatory regions in the inactive state of LRRK1. Disrupting the interface either results in kinase hyper-activation, possibly through lifting inhibitory Roc/COR-B/kinase interactions, or kinase inactivation, by preventing the formation of interactions required to stabilize the active state of the kinase, analogous to the R1305 mutation in the kinase αC helix.

**Figure 5:**
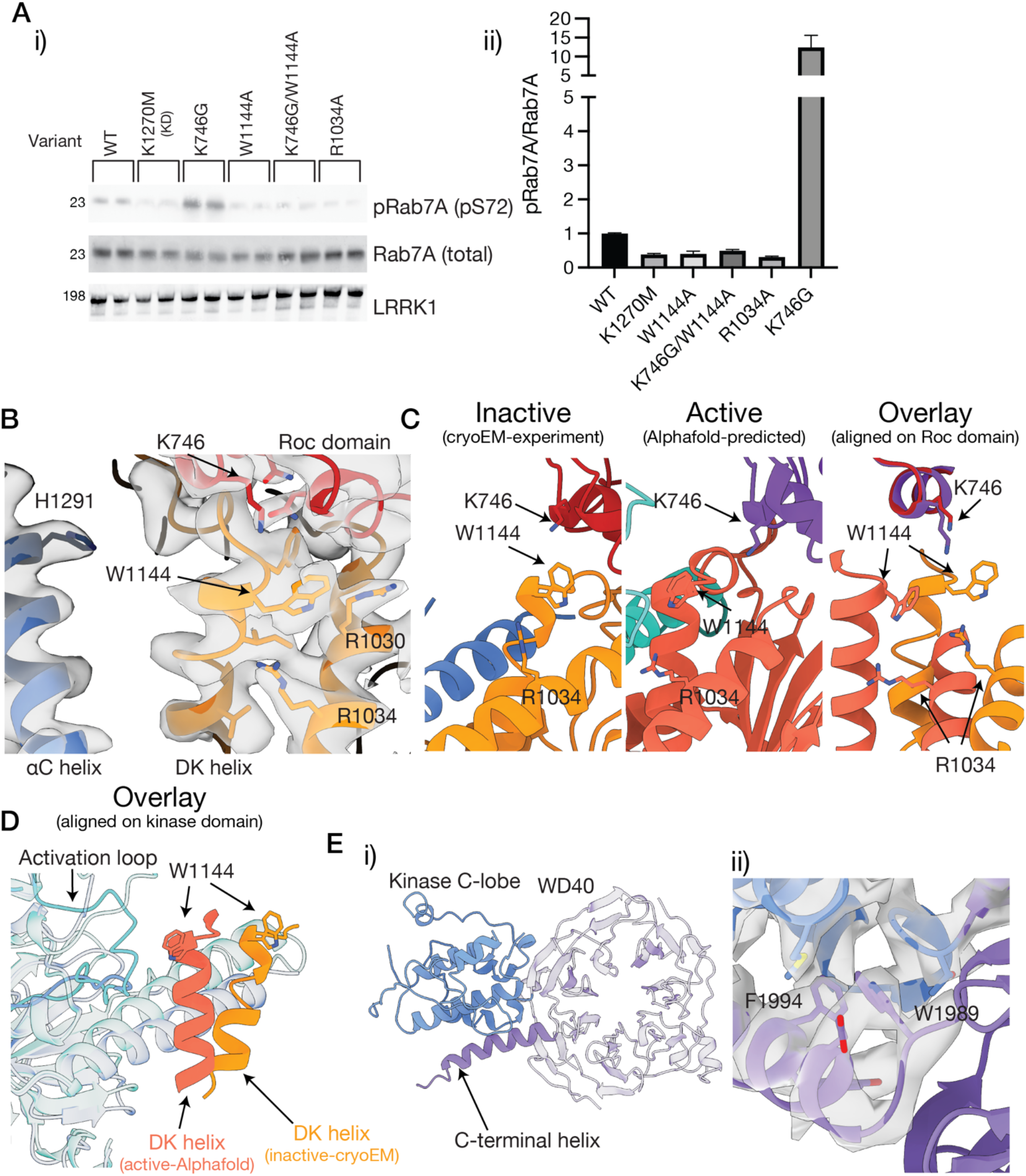
The Roc-COR-kinase interface and the kinase-WD40 interface. A) Effect of mutations in this interface on Rab7A phosphorylation by recombinant LRRK1, i) Western blot showing Rab7 phosphorylation, ii) quantification of multiple kinase assays from at least three independent protein preparations (see Methods, Supplementary Figure 7B for complete membrane images, values are mean ± SEM). B) Cryo-EM density supporting the Roc/COR interface, additionally showing that the αC helix is positioned away from the interface in the inactive state. C) Structural changes between the inactive-state cryo-EM model, and the activestate Alphafold model, highlighting a change in the position of K746 relative to the DK helix, and in the position of R1034. D) Overlay of the DK helix/activation loop interface between the cryo-EM and Alphafold models, highlighting the movement of the DK helix towards the activation loop. E) The interface formed by the LRRK1 kinase C-lobe and the WD40 repeats, and the additional helix in the kinase C-lobe, contributed by the C-terminal helix, i) the position of the helix, ii) cryo-EM density supporting the position of W1989 at the base of the helix, the deletion of which causes OSMD.

Finally, we showed that recombinant Y971F LRRK1, which is a known LRRK1 activating mutation *in celluo*^7,9,13^, is more active compared to WT LRRK1 in our *in vitro* assay (Supplementary Figure 8A, p=0.0021 compared to WT). Phosphorylation of LRRK1 at Y971 by EGFR results in kinase inhibition^13^, so the Y971F mutation has been proposed to activate LRRK1 by abolishing EGFR-mediated inactivation, although the Y971F mutation is activating in the absence of EGF simulation^7^, and is likewise activating in our *in vitro* assay. Y971 is located away from the kinase domain in the cryo-EM model (Supplementary Figure 8B), although it is located relatively close to the Roc domain and forms part of a hydrogen bonding network involving the Switch-I motif, which responds to guanosine nucleotide binding, so the effect of the Y971F mutation may be to indirectly modulate kinase activity through the LRRK1 Roc domain (Supplementary Figure 8C).

We also measured the effect of all mutations on the thermal stability of LRRK1 using differential scanning fluorimetry. All mutations modestly decreased the thermal stability by 1-2 °C (Supplementary Figure 9A-B), apart from the M1298A and Y971F mutations, which did not alter the thermal stability by more than 0.5 °C, and the D1409A kinase-dead mutation, which increased the stability by 0.9 °C. The K746G and K746G/W1144A mutations both had the greatest effect on the stability of LRRK1, reducing it by 2.5 °C and 2.0 °C respectively.

### The kinase domain interacts with the WD40 domain at the C-terminal helix

Analogous to LRRK2, the WD40 domain sits against the C-lobe of the kinase, with the WD40 C-terminal helix effectively contributing an additional helix to the kinase C-lobe fold^30^ (Figure 5Ei). The N-terminal end of the helix forming numerous interactions with the kinase C-lobe, largely through the aromatic residues W1989, F1997, Y1998 and Y2001 (Figure 5Eii). The helix is slightly bent, so the C-terminal end of the helix does not contact the C-lobe of the kinase. The exact role for the C-terminal helix is unclear. In LRRK2, the helix serves to bridge the kinase, COR and ankyrin domains^31^, and deletion of the C-terminal helix in LRRK2 results in insoluble protein^30^. In LRRK1, the C-terminal helix does not interact with either the COR or ankyrin domain. A two residue deletion (ΔW1989/ΔG1990) in the helix in LRRK1 causes OSMD^16^. These two residues sit at the ‘base’ of the C-terminal helix (Figure 5Eii). We attempted to express this mutant for biochemical characterization, however we did not observe any expression, showing that even subtle disruption of the C-terminal helix is not tolerated.

### The LRRs of LRRK1 are highly dynamic

Consensus refinements of LRRK1 resulted in a map with a significantly lower local resolution for the LRR domain, suggesting that the LRR is dynamic with respect to the remainder of the protein. Recent advances in cryo-EM data processing allow the interrogation of continuous motion from single-particle cryo-EM data^45,46^. To examine the structural dynamics we applied 3D variability analysis^46^ in *Cryosparc* to the cryo-EM dataset (Figure 6, Supplementary Movie), solving for three variability modes. Briefly, each variability mode corresponds to a vector which correspond to trajectories in the space in which the molecule experiences variability. The first variability mode corresponds to little molecular motion. The second and third variability components corresponds to domain movements, pivoting of the Roc/COR domain/kinase N-lobe relative to the kinase C-lobe and WD40 domain, along with a correlated large movement of the LRRs (Figure 6). Similar flexibility in the Roc-COR domain was noted in the low-resolution cryo-EM map of the C-terminal domain of LRRK1^32^. These structural dynamics in the kinase/COR domains in LRRK1 may reflect inter-domain transitions between active and inactive conformations of the COR and kinase domain. The dynamics in the LRR domain may serve to sterically control access to the kinase, with configurations open for substrate binding and the more occluded position sterically hindering substrate binding. Presumably, only the relatively open configurations would allow for substrate binding.

**Figure 6:**
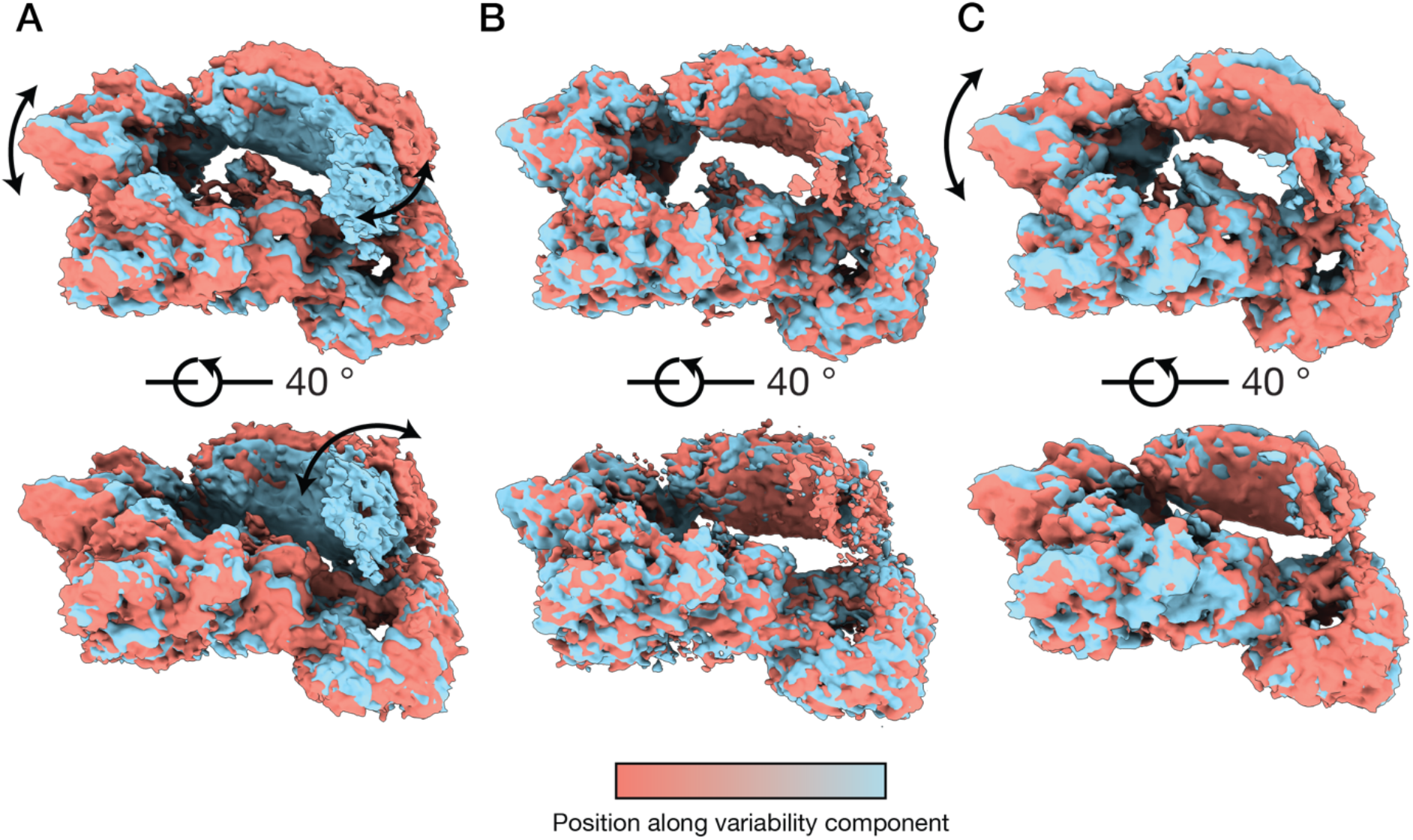
3D variability analysis of the LRRK1 monomer. A) variability component 1, showing a movement of the Roc domain, and a large movement of the leucine-rich repeats. B) variability component 2, showing little movement. C) variability component 3, showing a movement of the Roc domain. The arrows indicate the movement experienced by LRRK1 along the variability component, with the density maps colored according to their position along the variability component. See also, Supplementary Movie.

### Conclusion

Here, we have presented the structure of the full-length LRRK1 monomer, alongside a lower-resolution cryo-EM map of the LRRK1 dimer. The structures of the LRRK1 monomer and dimer are striking in their differences to recent experimentally determined structures of LRRK2, particularly in the position of the LRRs in the monomer. Additionally, we show an intrinsically inactive form of the LRRK1 dimer, in which both kinase domains are not only in an inactive conformation but also occluded for substrate binding by the ankyrin repeats from the neighboring molecule. Activation of this dimer would probably require dissociation of the dimer to monomers. It should be noted that our results show the full-length inactive LRRK1 exists as both monomer and dimer in solution, suggesting the inactivation-activation cycle of the LRRK1 monomer can be an independent event, which does not necessarily to be a subsequent downstream step of dimer-to monomer transition, with further work required to determine what controls the LRRK1 dimer-to-monomer transition. Additionally, it will be important to determine the physiological relevance of various oligomerization states obtained under cryo-EM conditions for both LRRK1 and LRRK2.

The structure and biochemical studies highlight the kinase and GTPase interdomain contacts which likely serve to regulate the activity of the LRRK1 kinase. In particular, the extended αC helix, which is unique to LRRK1, serves as a key regulatory scaffold, with mutations in the helix serving to inactivate the kinase through disrupting interactions formed in the kinase active state. The extended αC helix interaction with the COR-B domain likewise serves to stabilize the inactive state of the kinase, disrupting this interface serves to activate the kinase. More distant residues, located at the Roc/COR and in the COR domain more cryptically serve to modulate kinase activity, hinting at the large structural rearrangements, especially involving the DK helix, that must occur to enable LRRK1 kinase activation. Fully understanding this activation mechanism will require experimentally determined structures of LRRK1 in the active state, in particular phosphorylated by its regulatory partners such as PKC. Overall, these results provide a framework for future studies examining LRRK1 biology and enable a comparison of the mechanisms underpinning control of both LRRKs in normal physiology and disease.

## Methods

### Expression and purification of LRRK1

We purchased the full-length human LRRK1 gene from Genscript (residues 1-2015, uniprot Q38SD2), codon optimized for *Homo sapiens.* We sub-cloned the codon-optimized LRRK1 sequence into the pFastBac vector for insect cell protein expression, N-terminally fused to a 3xFlag affinity tag and rhinovirus 3C protease cleavage site, using the NEB HiFi DNA assembly kit (cat. E2621S). We generated bacmid using the Bac-to-Bac system (Invitrogen) and transfected the bacmid into Sf9 cells to generate a baculovirus stock. We subsequently generated a high titer baculovirus stock for protein expression.

4 L exponentially growing Sf9 cells at a density of 2-3 × 10^6^ cells/mL in Sf900-III SFM media (Invitrogen) was infected with the high-titer baculovirus stock. Cells were incubated for 48-72 hours at 27 °C with shaking post-infection, and then harvested by centrifugation. The cell pellet was resuspended in resuspension buffer [20 mM Tris, 10 mM CaCl_2_, 5 mM MgCl_2_, 100 mM NH_4_Cl, 100 mM NaCl, 50 mM L-Arg, 50 mM L-Glu, 0.0008% Tween-80, 10% glycerol pH 8.3]. Following resuspension, protease inhibitors (Roche), 10 mM β-glycerophosphate and 1 mM sodium vanadate (to inhibit phosphatases) were added to the pellet, and the pellet was frozen at −80 °C.

All purification steps were done at 4 °C. For purification, the cell pellet was thawed, and cells were lysed using multiple passes through a fluidizer (LM20 Microfluidizer, Microfluidics Corp). The crude lysate was centrifuged, and the supernatant taken and applied to a Flag-M2 affinity column (Sigma cat. A2220) and incubated for 2-3 hours. The column was then washed once with resuspension buffer, twice with wash buffer [20 mM Tris, 500 mM NaCl, 5 mM MgCl_2_, 0.0008% Tween-80 pH 8.3], and then once with gel filtration buffer [20 mM Tris, 150 mM NaCl, 5 mM MgCl_2_, 0.0008% Tween-80 pH 8.3]. For elution, the column was washed three times with gel filtration buffer supplemented with 150 μg/mL 3x-Flag peptide (Genscript cat. RP21087, Supplementary Figure 1A).

Eluted fractions containing LRRK1 were concentrated to ~500 μL using a centrifugal concentrator (100 kDa cutoff, Pall cat. MAP100C38) and applied to a Superose 6 Increase 10/30 column (GE Healthcare), equilibrated in gel filtration buffer. The fraction at ~15.5 mL was used for LRRK1 monomer grid preparation (see Supplementary Figure 1). The fraction at ~14.5 mL was used for LRRK1 dimer grid preparation. Protein was concentrated and used immediately for cryo-EM grid preparation and biochemical/biophysical characterization. The concentration was assessed using UV spectroscopy, using an extinction coefficient of 1.1 cm^-1^(mg/mL)^-1^.

For small-scale affinity purification for kinase assays, mutations were introduced into the full-length, codon-optimized LRRK1 gene using the NEB Q5 Site-Directed Mutagenesis Kit (cat. E0554), and baculovirus was generated as above. Small-scale expressions were done in a volume of 50-250 mL Sf9 cells in Sf900-III media, at a density of 2 × 10^6^ cells/mL. Cells were infected with baculoviruses, incubated for 60-72 hours at 27 °C with shaking post-infection, and then harvested by centrifugation. The cell pellet was frozen once, and then resuspended in lysis buffer [20 mM Tris, 10 mM CaCl_2_, 5 mM MgCl_2_, 100 mM NH_4_Cl, 100 mM NaCl, 50 mM L-Arg, 50 mM L-Glu, 1% Tween-80, 20 μM GppNHP, 10% glycerol pH 8.3], supplemented with protease inhibitors (Roche), 10 mM β-glycerophosphate and 1 mM sodium vanadate (to inhibit phosphatases), and incubated for one hour at 4 °C. Following lysis, the lysate was clarified by centrifugation, and the clarified lysate added to 50 μL Flag-M2 affinity resin (Sigma cat. A2220) and incubated at 4 °C for two hours in 1.5 mL Eppendorf tube. The resin was washed as above, with the exception that 20 μM GppNHP was added to all buffers, and then eluted with a single wash of gel filtration buffer supplemented with 150 μg/mL 3x-Flag peptide (Genscript cat. RP21087) and 20 μM GppNHP. The eluate was filtered using a 0.22 μm Ultrafree-GV centrifugal filter (Sigma cat. UFC30GVNB) to remove any free resin, and then used in a kinase assay. Protein concentration was assessed using UV spectroscopy, using an extinction coefficient of 1.1 cm^-1^(mg/mL)^-1^.

### Mass photometry

Mass photometry data was collected using a OneMP mass photometer (Refeyn). 15 μL of detergent-free gel filtration buffer [20 mM Tris, 150 mM NaCl, 5 mM MgCl_2_ pH 8.3] was applied to a coverslip, and, after focusing, 3 μL of the undiluted gel filtration fraction was added to the drop and mixed. Movies were acquired for 6,000 frames (60 seconds) using AcquireMP software with the large view setting. Raw data was processing using DiscoverMP software. For calibration, a mixture of beta amylase (Sigma cat. A8781) and thyroglobulin (Sigma cat. T9145) was used.

### Multi-angle light scattering

SEC-MALS data were collected using a Shimadzu LC-20AD HPLC, coupled to a Shimadzu SPD-20A UV detector, Wyatt Dawn MALS detector and Wyatt Optilab refractive index detector. Data were collected following in-line fractionation with a Superose 6 15/150 column (GE Healthcare), pre-equilibrated in gel filtration buffer, running at a flow rate of 0.3 mL/min. 50 μL of the monomer peak from gel filtration chromatography were applied to the column for analysis. Data were analyzed using ASTRA v. 8.0.2.5 (Wyatt). Detector response was normalized using monomeric BSA (Thermo Fisher, cat. 23209). Protein concentration was determined using differential refractive index, using a dn/dc of 0.184.

### Expression and purification of Rab7A

The sequence for Rab7A (residues 2-176, uniprot P51149) was purchased from Genscript, N-terminally fused to a His tag and tobacco etch virus (TEV) protease cleavage site, codon-optimized, and subcloned into a pET29a vector for expression in *E. coli.* The construct was transformed into sHuffle T7 Express cells^47^ (New England Biolabs, cat. C3029J) for protein expression. For protein expression, 6 L of transformed cells in lysogeny broth (LB), supplemented with 50 μg/mL kanamycin, were grown at 37 °C until OD600 ~0.6-0.7, cultures were induced with 1 mM IPTG and grown for 16-18 hours at 16 °C, cultures were subsequently harvested by centrifugation, and resuspended in wash buffer [20 mM Tris, pH 7.4, 500 mM NaCl, 20 mM imidazole, 10 mM MgCl_2_]. Protease inhibitors (Roche) were added to the pellet following harvest, and the pellet was frozen at −80 °C.

All purification steps were done at 4 °C. For purification, the cell pellet was thawed, and cells were lysed using multiple passes through a fluidizer (LM20 Microfluidizer, Microfluidics Corp). The crude lysate was centrifuged, benzonase was added to the lysate (Sigma cat. E1014) and incubated for 15 minutes at room temperature. The cell lysate was then 0.45 μm filtered and applied to a 5 mL HisTrap HP column (GE Healthcare). The column was washed with five column volumes of wash buffer, then eluted with elution buffer [20 mM Tris, pH 7.4, 500 mM NaCl, 500 mM imidazole, 10 mM MgCl_2_] over a gradient of twenty column volumes. Fractions containing Rab7A were then pooled. TEV protease was added to the pooled protein (prepared in-house), and then dialyzed overnight against gel filtration buffer [20 mM Tris, pH 7.4, 150 mM NaCl, 10 mM MgCl_2_]. The following day, the dialysate was taken, and the imidazole and salt concentration adjusted to 20 mM and 500 mM, respectively, and applied to a 5 mL HisTrap HP column to remove TEV protease, free His tags, and uncleaved Rab7A. The flowthrough was taken, which contained Rab7A without a His tag, concentrated to ~5 mL and applied to a Superdex 75 16/160 column (GE Healthcare), equilibrated in gel filtration buffer. Fractions containing purified Rab7A were taken, pooled, concentrated to ~10 mg/mL and frozen at −80 °C for long-term storage. Typical yields were 1-3 mg/L cell culture, with protein concentrations determined using extinction coefficients calculated from the protein sequence.

### LRRK1 kinase activity assays

LRRK1 kinase assays with purified LRRK1 monomer were set up in a 15 μL final mixture with 150 nM LRRK1 monomer, 10 μM Rab7A, 5 mM ATP and 2 mM GppNHP, in 20 mM Tris, 150 mM NaCl, 5 mM MgCl_2_ and 0.0008% Tween-80. The kinase reaction was carried out at 30 °C for two hours with shaking (300 rpm) in a Thermomixer (Eppendorf). The reaction was stopped by addition of 5 μL of 4X SDS-PAGE loading buffer [250 mM Tris, 8% SDS, 0.2% bromophenol blue, 40% glycerol, 20% β-mercaptoethanol], heated for 10 minutes at 95 °C, and the samples frozen (−80 °C) prior to immunoblot analysis.

LRRK1 kinase assays with affinity-purified LRRK1 and LRRK1 mutants were set up in 30 μL final mixture with 100 nM LRRK1, 1 μM Rab7A, 5 mM ATP and 2 mM GppNHP, in 20 mM Tris, 150 mM NaCl, 5 mM MgCl_2_ and 0.0008% Tween-80. The kinase reaction was carried out at 30 °C for thirty minutes with shaking (300 rpm) in a Thermomixer (Eppendorf). The reaction was stopped by addition of 5 μL of 4X SDS-PAGE loading buffer [250 mM Tris, 8% SDS, 0.2% bromophenol blue, 40% glycerol, 20% β-mercaptoethanol], heated for 10 minutes at 95 °C, and the samples frozen (−80 °C) prior to immunoblot analysis. Assays were performed from a total of three independent protein preparations in duplicate (for a total of six independent kinase assays, eighteen assays for the WT and K1270M kinase-dead mutants).

Subsequently, samples were resolved on 4-12% Bis-Tris gradient gels. Protein was subsequently wet-transferred to a 0.4 μm PVDF membrane. Membranes were subsequently blocked for 30 minutes using 5% bovine serum albumin (BSA) in TBS-T [20 mM Tris, 150 mM NaCl pH 7.4, 1% Tween-20] and probed with anti-phospho Rab7A primary antibody (Abcam cat. ab302494, diluted 1:1000), Rab7A (total) primary antibody (Abcam cat. ab50533, diluted 1:2000) and anti-LRRK1 primary antibody (Abcam cat. ab228666, diluted 1:2000). Membranes were washed using TBS-T and then probed with goat anti-rabbit IR-fluorescent secondary antibody (LiCor cat. 926-3221, diluted 1:20,000) and goat anti-mouse IR-fluorescent secondary antibody (LiCor cat. 926-68072). The membrane was subsequently washed with TBS-T and imaged using a Typhoon scanner (GE Healthcare). Blots were quantified using Fiji^48^ to determine the pRab7A/Rab7A ratio, this ratio was then normalized to LRRK1 WT reactions on the same membrane. Significance of differences was quantified using a one-way Brown-Forsythe and Welch ANOVA test in Prism 9.4.1, using an unpaired t-test with Welch’s correction to compare mutants with the WT and K1270M kinase-dead mutant.

### Cryo-EM – grid preparation, data collection and 3D reconstruction for the LRRK1 monomer

Cryo-EM was performed at the cryo-EM facility in the Center for Structural Biology, NCI-Frederick. 1.5 μL purified LRRK1 monomer or dimer at a concentration of 0.2-0.5 mg/mL was applied to each side of a Quantifoil R 1.2/1.3 Gold 300 Mesh grids (cat. Q3100AR1.3), that had been glow discharged on each side (25 mA for 30 sec). Grids were vitrified using a Leica EM GP2 plunge freezer, with a blotting time of 1-3 seconds. Grids were subsequently imaged using a Gatan K3 direct detector, equipped with an energy filter, mounted on a Talos Arctica G2 (Thermo Fisher) electron microscope in super resolution mode (pixel size 0.405 Å/pixel, 100,000 × magnification). 50 frames per movie were acquired for a total dose of approximately 50 elections/Å^-2^. Data was collected using the EPU program (Thermo Fisher), with defocus values ranging from −2.5 to −0.8 μm. 22,865 movies were collected from LRRK1 monomer grids, and 10,021 movies from LRRK1 dimer grids, for a total of 32,886 movies.

Data processing was performed using Cryosparc 3.3^49^. Movies were imported into Cryosparc, patch-motion and patch-CTF corrected. Movies were binned to the physical pixel size in the patch motion step. Movies with a CTF resolution >5 Å were removed from the stack, leaving 18,074 monomer movies and 8,612 dimer movies for further processing (26,686 movies total). 3,553,719 particles were picked from the monomer movies and 1,133,944 from the dimer movies (for a total of 4,687,663 particles), using a Topaz model trained on an initial LRRK1 monomer or dimer dataset^50,51^. Particle stacks were subsequently merged and curated using two rounds of 2D classification to remove clear false positive particles, carbon edges and junk particles, with duplicate particles which were introduced due to re-centering removed after each round of 2D classification. The 517,408 particles were used in a five-class *ab initio* model generation in Cryosparc. Separately, 26,352 particles corresponding to the LRRK1 dimer were used to generate an initial model of the LRRK1 dimer. These models were used as the input for a seven-class heterogenous refinement with the full stack of 517,408 particles (Supplementary Figure 2). Particles from the dimer class was used in a subsequent non-uniform refinement^52^ step, giving a map with an overall resolution of 6.38 Å, judged by the gold standard FSC in Cryosparc (Supplementary Figure 2). Particles from monomer classes were used in a second heterogenous refinement round, with four classes, with the 183,273 particles from the best class used in a final non-uniform refinement step, giving a map with an overall resolution of 3.92 Å, judged by the gold standard FSC in Cryosparc (Supplementary Figure 2). Subsequently, 3D variability analysis^46^ was used in Cryosparc to analyze the structural dynamics of LRRK1. Three variability modes were solved, and 3D variability was visualized using ‘intermediates’ mode, with ten frames and a filter resolution of 4 Å.

Inspection of the map revealed that the LRR region was poorly defined relative to the rest of the map. To improve the resolution of the C-terminal region, the LRR region was subtracted from the monomer map using Particle Subtraction in Cryosparc. Local refinement in Cryosparc, using a soft mask corresponding to the C-terminal region, was then used to improve the resolution of the C-terminal region to 3.78 Å. Local resolution maps and FSC curves were generated using Cryosparc. For a graphical summary of the cryo-EM image processing, see Supplementary Figure 3. For a summary of cryo-EM reconstruction statistics, see Supplementary Table 1. We combined the Cryosparc-sharpened global and local refinement maps of the monomer using the *phenix.combine_focused_maps* in Phenix 1.20^53^ for atomic model refinement.

### Cryo-EM – atomic model refinement

For atomic model building and refinement, we first pre-processed the Alphafold^41,54^ predicted model of LRRK1 (AF-Q38SD2-F1) using *phenix.process_predicted_model* in Phenix 1.20^53^, which removed low-confidence regions in the predicted model, and split the model into domains for subsequent rigid body fitting. We fit the domains into the map using *UCSF Chimera*^55^ to generate an initial model. We refined this model against the composite cryo-EM map using *phenix.real_space_refine*^56^, followed by manual model building in *Coot*^57^ and subsequent refinement in *phenix.real_space_refine.* Geometry and real-space correlation validation was performed using the *phenix.validation_cryo-EM* tool (incorporating *MOLProbity*^58^). Figures were prepared using *UCSF ChimeraX*^59,60^. For a summary of model building and validation statistics, see Supplementary Table 1.

For the LRRK1 dimer atomic model, two copies of the LRRK1 monomer were docked into the map using *UCSF Chimera,* and the ankyrin repeats from the Alphafold model of LRRK1 were added. This model was subsequently refined using *phenix.real_space_refine* using five rounds of rigid body refinement, defining the ANK repeats, LRRs, Roc-COR and kinase-WD40 domains as separate rigid bodies. Due to the low resolution of the map, we did not deposit an atomic model in the PDB.

### Differential scanning fluorometry

Thermal stability was measured using the Prometheus NT.48 nano-DSF instrument (NanoTemper). LRRK1 or the relevant mutant in the buffer 20 mM Tris, 150 mM NaCl, 5 mM MgCl_2_, 0.0008% Tween-80 pH 8.3, at a concentration of 0.1 mg/mL was loaded into a nanoDSF glass capillary. Thermal unfolding was measured at a heating rate at 1 °C, protein melting temperature was calculated from the first derivative of the ratio of tryptophan fluorescence at 330 nm and 350 nm.

## Acknowledgements

We thank Dr. Dan Shi for assistance collecting the cryo-EM data, Dr. Sergey G. Tarasov and Marzena Dyba for assistance with collecting the differential scanning fluorometry and mass photometry data. Cryo-EM data was collected using the Talos Arctica G2 in the Center for Structural Biology cryo-EM facility, NCI at Frederick. We acknowledge use of the Biophysics Resource, Center for Structural Biology, NCI at Frederick, and use of the Frederick Research Computing Environment cluster. This research was supported by federal funds from the intramural program of the National Cancer Institute, National Institutes of Health, under project number ZIA BC 011744 (P.Z.).

## Author Contributions

R.M. and J.A.M.F. collected cryo-EM data, R.M. processed the cryo-EM data with input from J.A.M.F. and P.Z, R.M. prepared all samples and undertook the biochemical/biophysical characterization. P.Z. supervised the work. R.M. wrote the first draft of the manuscript, R.M., J.A.M.F. and P.Z. edited the manuscript. All authors approved the current version of the manuscript.

## Competing Interests

No competing interests.

## Data and materials availability

All data needed to evaluate the conclusions in the manuscript are present in the manuscript and/or the Supplementary Materials. Maps have been deposited in the EMDB with accession codes EMD-28950 (LRRK1 monomer composite map), EMD-28949 (LRRK1 monomer global refinement), EMD-28951 (LRRK1 C-terminus local refinement after subtraction of the LRRs), and EMD-28952 (LRRK1 dimer). The atomic model of the LRRK1 monomer has been deposited in the PDB with accession code 8FAC.

## Notes

### Competing Interest Statement

The authors have declared no competing interest.

