## Supplementary Material for "Structure and regulation of full-length human leucine-rich repeat kinase 1"

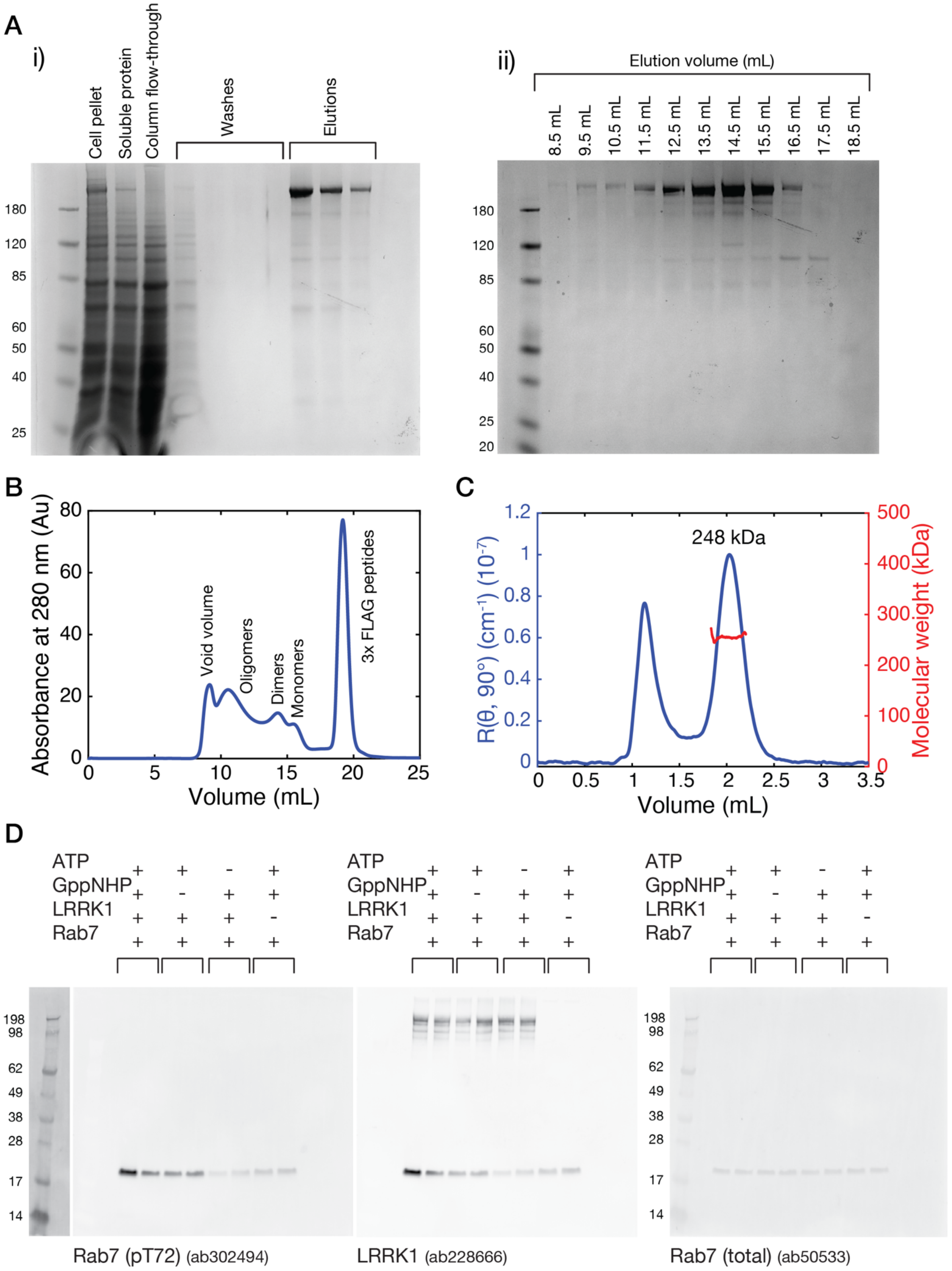

A
i)
Cell pellet
Soluble protein
Column flow-through
Washes
Elutions
180
120
85
60
50
40
25
ii)
Elution volume (mL)
8.5 mL
9.5 mL
10.5 mL
11.5 mL
12.5 mL
13.5 mL
14.5 mL
15.5 mL
16.5 mL
17.5 mL
18.5 mL
180
120
85
60
50
40
25
20
B
Absorbance at 280 nm (Au)
Void volume
Oligomers
Dimers
Monomers
3x FLAG peptides
Volume (mL)
C
248 kDa
R(θ, 90°) (cm⁻¹) (10⁻⁷)
Molecular weight (kDa)
Volume (mL)
D
ATP
GppNHP
LRRK1
Rab7
198
98
62
49
38
28
17
14
Rab7 (pT72) (ab302494)
LRRK1 (ab228666)
Rab7 (total) (ab50533)

**Supplementary Figure 1:** Purification of LRRK1. A) 4-12% SDS-PAGE gels showing LRRK1 purity following, i) Flag-affinity purification and ii) gel filtration chromatography on a Superose 6 Increase 10/30 gel filtration chromatography column. B) Gel filtration chromatogram following Flag-affinity chromatography. Peaks corresponding to the void volume, high-molecular weight oligomers, dimers and monomers are indicated. C) SEC-MALS chromatogram for the LRRK1 monomer. D) Complete membrane images for the Western blots shown in Figure 1C.

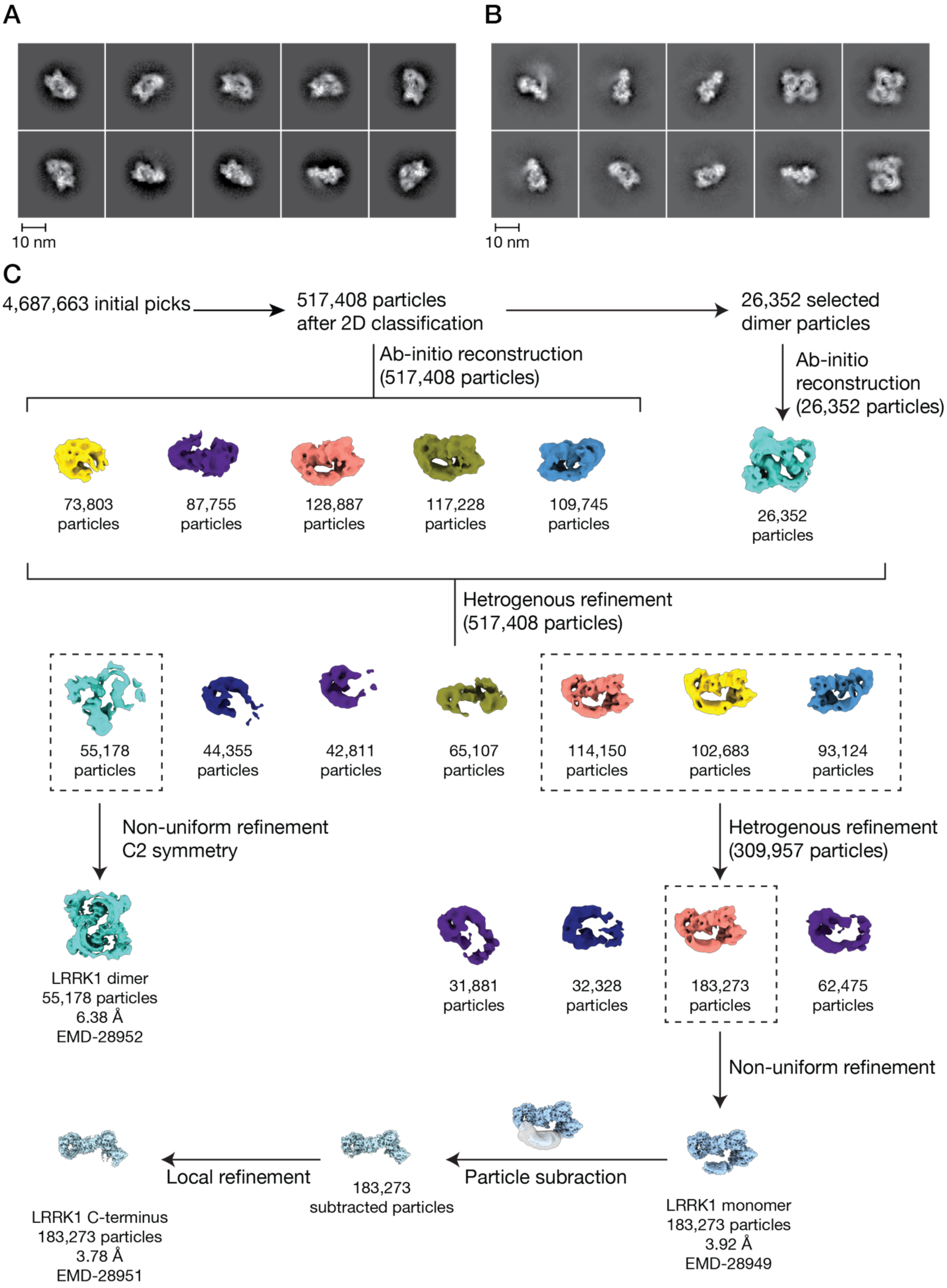
A
B
10 nm
10 nm
C
4,687,663 initial picks
517,408 particles after 2D classification
26,352 selected dimer particles
Ab-initio reconstruction (517,408 particles)
Ab-initio reconstruction (26,352 particles)
73,803 particles
87,755 particles
128,887 particles
117,228 particles
109,745 particles
26,352 particles
Hetrogenous refinement (517,408 particles)
55,178 particles
44,355 particles
42,811 particles
65,107 particles
114,150 particles
102,683 particles
93,124 particles
Non-uniform refinement C2 symmetry
Hetrogenous refinement (309,957 particles)
LRRK1 dimer
55,178 particles
6.38 Å
EMD-28952
31,881 particles
32,328 particles
183,273 particles
62,475 particles
Non-uniform refinement
Local refinement
Particle subraction
183,273 subtracted particles
LRRK1 C-terminus
183,273 particles
3.78 Å
EMD-28951
LRRK1 monomer
183,273 particles
3.92 Å
EMD-28949

**Supplementary Figure 2:** Cryo-EM data collection and processing for the LRRK1 monomer. A) Representative 2D class averages for data collected from LRRK1 monomer grids. B) Representative 2D class averages for data collected from LRRK1 dimer grids. C) Summary of cryo-EM 3D reconstruction and refinement.

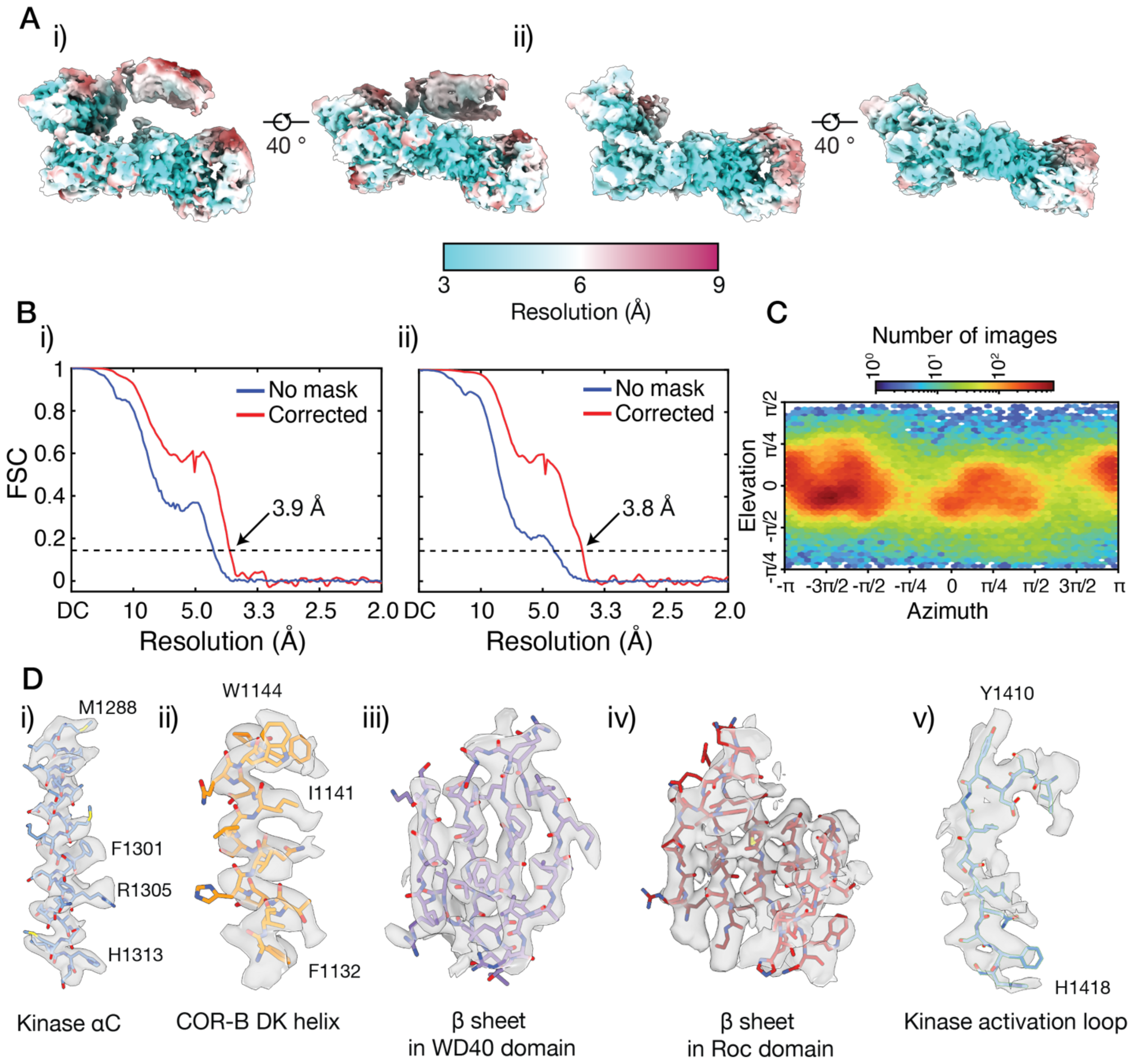

**Supplementary Figure 3:** Cryo-EM resolution estimation for the LRRK1 monomer. A) Local resolution maps for i) the global non-uniform refinement map, ii) the local refinement map for the RCKW domains. B) Gold-standard Fourier shell correlation (FSC) curves for, i) the global non-uniform refinement map, ii) the local refinement map. C) Viewing direction distribution for the global non-uniform refinement map. D) Representative density for the, i) kinase αC helix, ii) the COR-B DK helix, iii) a β-sheet in the WD40 domain, iv) a β sheet in the Roc domain, v) the kinase domain activation loop.

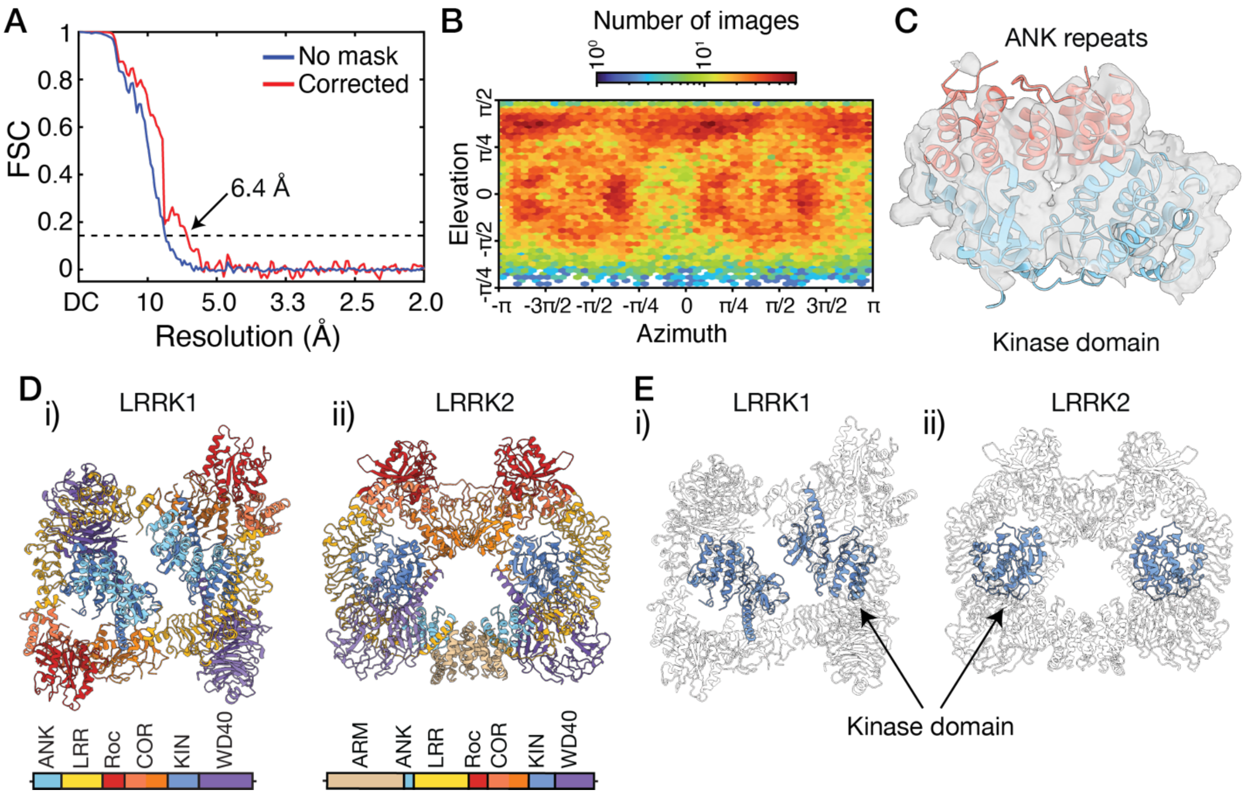

**Supplementary Figure 4:** Cryo-EM analysis of the LRRK1 dimer. A) Gold-standard FSC curve. B) Viewing direction distribution for the global non-uniform refinement map. C) Additional density observed above the kinase domain, assigned to the ankyrin repeats from the opposing molecule in the dimer. D) Comparison of the i), LRRK1 and ii), LRRK2 dimers. E) Comparison of the position of the kinase domain in the i), LRRK1 and ii), LRRK2 dimers.

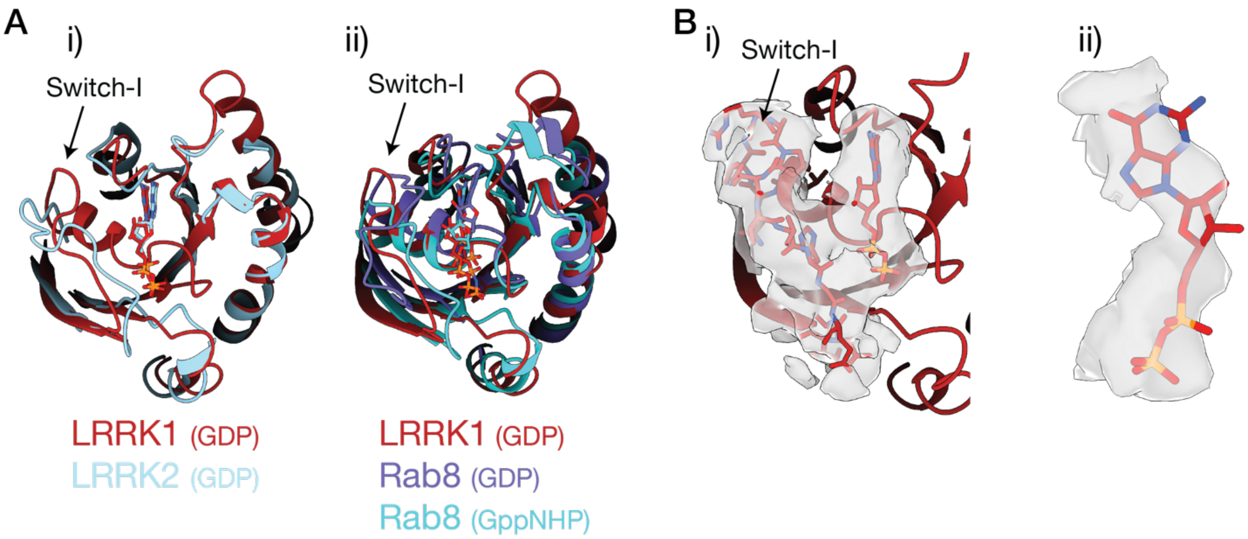

**Supplementary Figure 5:** The nucleotide state of the Roc domain. A) shows an overlay of the LRRK1 Roc domain with the Roc domain from i) LRRK2, bound to GDP (PDB: 7HLW[1]), ii) the small GTPase Rab8 bound to GDP (PDB: 4LHV[2]) and Rab8 bound to GppNHP (PDB: 4LHW), showing that the Switch-I motif in LRRK1 adopts a position consistent with GDP binding. B) i) Density supporting the position of GDP, the Switch-I motif and nearby residues, ii) Density supporting GDP.

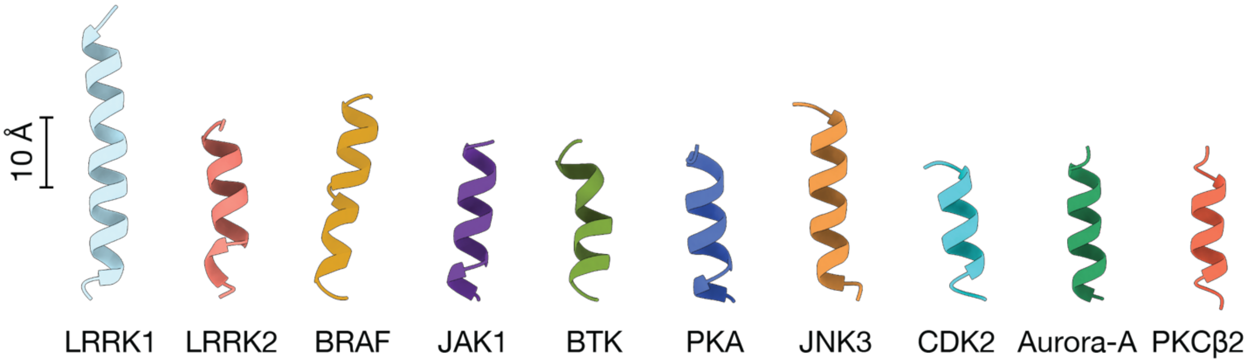

**Supplementary Figure 6:** Comparison of the αC helix from LRRK1 with other protein kinases. Alongside LRRK1, the αC helix from LRRK2 (PDB: 7LHW[1]), BRAF (PDB: 7MFD[3]), JAK1 (PDB: 6C7Y[4]), BTK (PDB: 1K2P[5]), PKA (PDB: 2CPK[6]), JNK3 (PDB: 1JNK[7]), CDK (PDB: 1FIN[8]), Aurora-A kinase (PDB: 1MQ4[9]) and PKCβ2 (PDB: 3PFQ[10]). Each αC helix is shown on the same scale, indicating that the LRRK1 helix is atypically long.

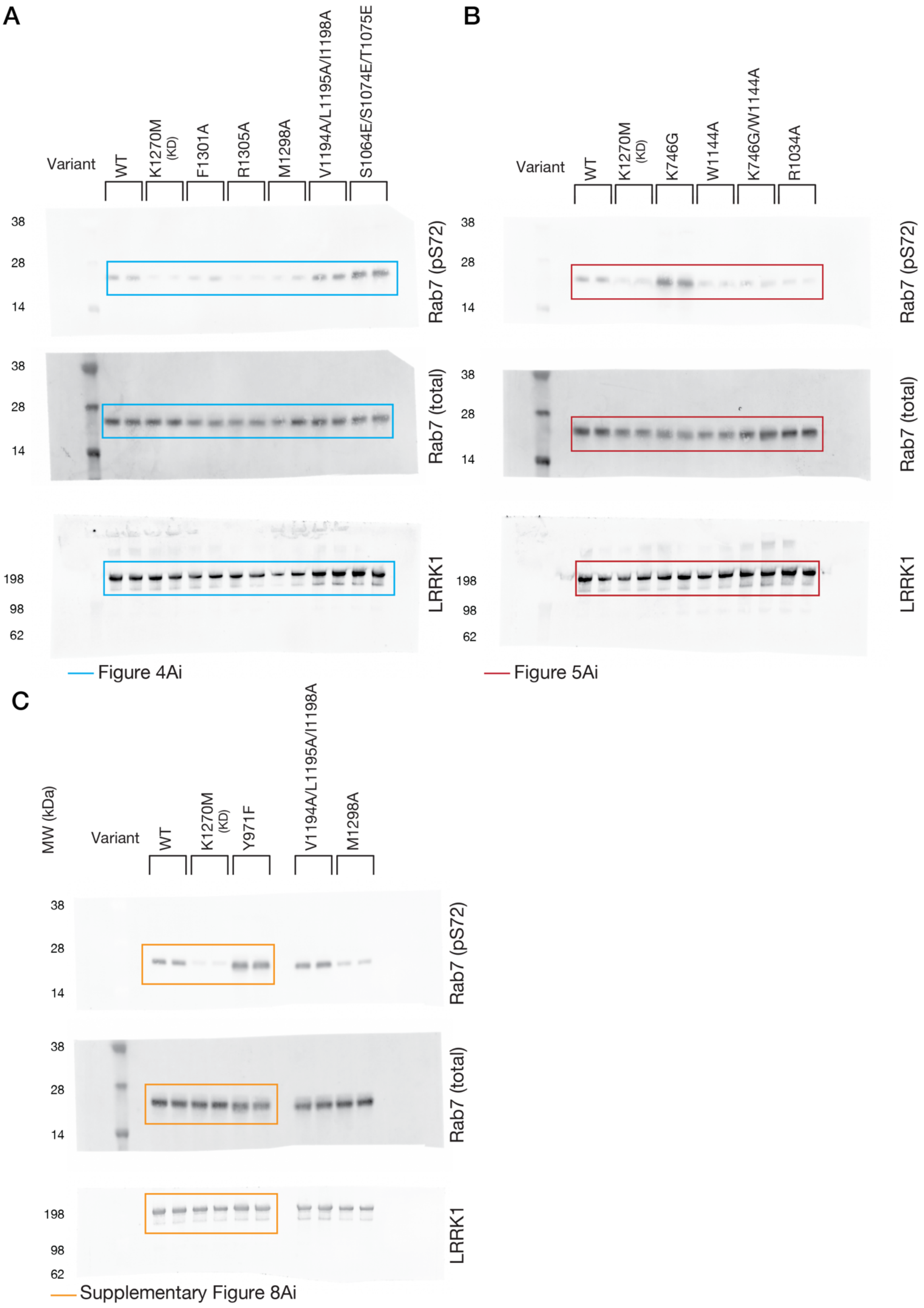
A
Variant
WT
K1270M (KD)
F1301A
R1305A
M1298A
V1194A/L1195A/I1198A
S1064E/S1074E/T1075E
38
28
14
Rab7 (pS72)
Rab7 (total)
198
98
62
LRRK1
Figure 4Ai
B
Variant
WT
K1270M (KD)
K746G
W1144A
K746G/W1144A
R1034A
Figure 5Ai
C
MW (kDa)
Variant
WT
K1270M (KD)
Y971F
V1194A/L1195A/I1198A
M1298A
Supplementary Figure 8Ai

**Supplementary Figure 7:** Complete membrane images for the membranes shown in Figure 4Ai, A), the membranes shown in Figure 5Ai B) and the membranes shown in Supplementary Figure 8Ai, C). Note membranes were cut to allowing simultaneous probing with multiple antibodies.

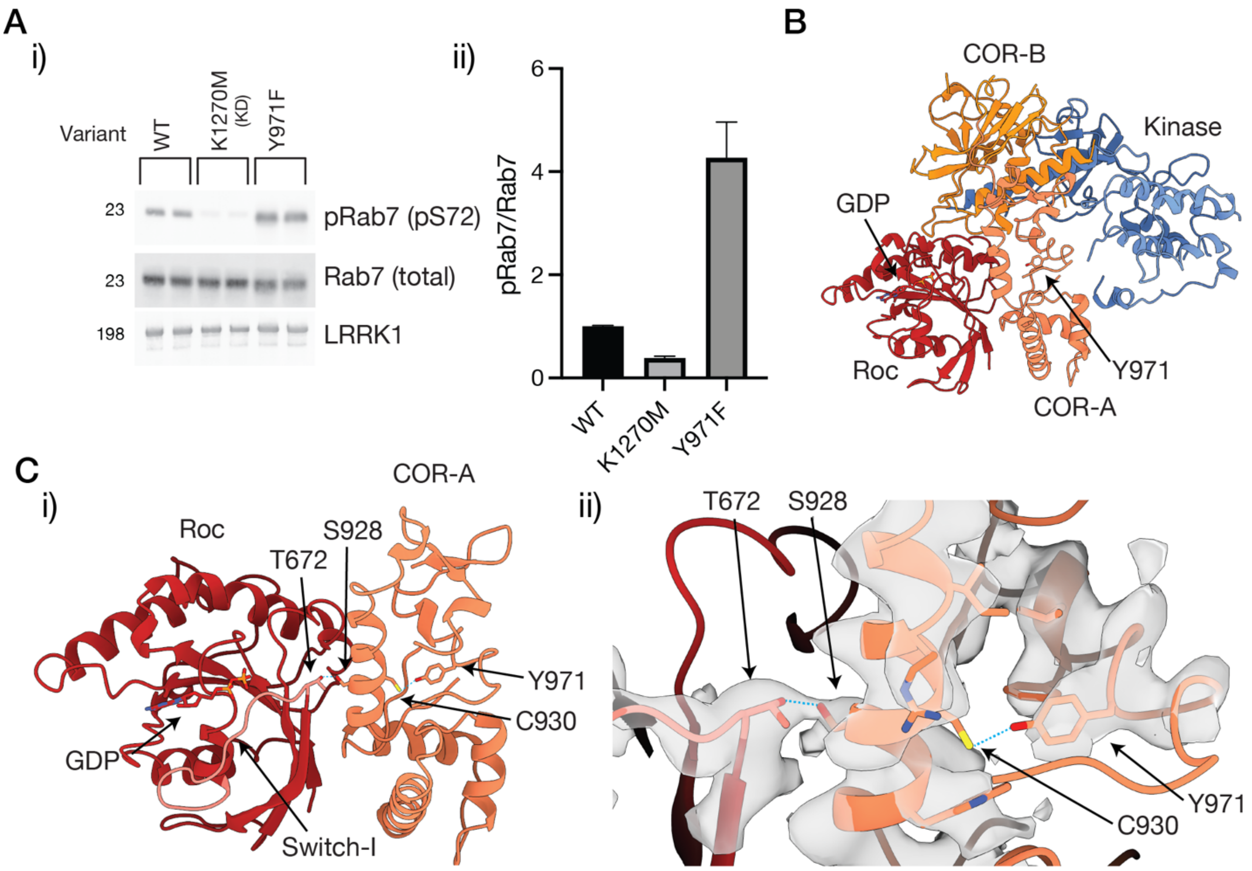

**Supplementary Figure 8:** Activation of LRRK1 by the Y971F mutation, and location in the structure. A) Western blot i), and quantification, ii) showing that recombinant Y971F LRRK1 increases Rab7 phosphorylation ~5 fold over WT LRRK1, error bars are SEM, see Supplementary Figure 7C for complete membrane images. B) Location of Y971 in the structure, in the COR-A domain. C) Position of Y971 in an interaction network involving the Switch-I motif in the GTPase domain (colored light red) i), cryo-EM density supporting the position of Y971 and other residues, ii).

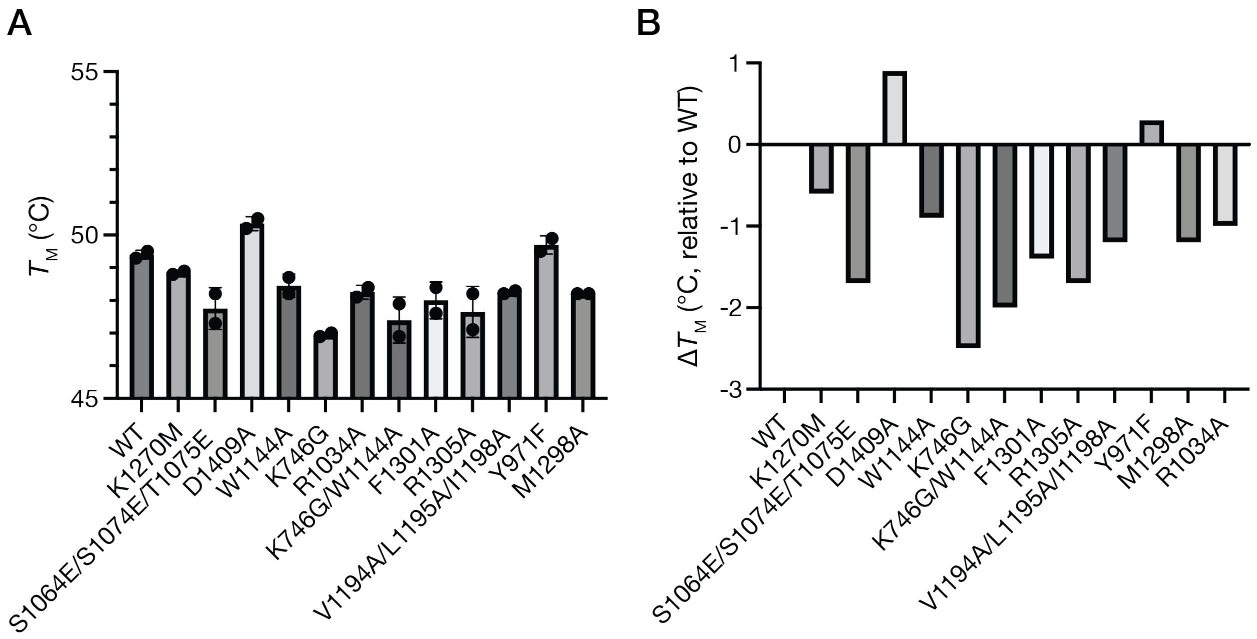

**Supplementary Figure 9:** Thermal stability of LRRK1 variants. A) Measured melting temperature ($T_M$) for fourteen LRRK1 variants, measured using a Nanotemper Nano-DSF instrument. B) Melting temperature relative to WT LRRK1. Each measurement was conducted using two separate protein preparations.

**Supplementary Table 1:** Cryo-EM data collection, 3D reconstruction and model building statistics.

| | LRRK1 monomer | LRRK1 C-terminus (LRRs subtracted, local refinement) | LRRK1 dimer |
|---|---|---|---|
| **Data collection and image processing** | | | |
| Microscope | Talos Arctica | | |
| Camera | K3 | | |
| Magnification | 100,000 | | |
| Electron energy (kV) | 200 | | |
| Electron exposure ($e^{-}A^{-2}$) | 50 | | |
| Defocus range (µm) | 0.8-2.5 | | |
| Physical pixel size | 0.81 | | |
| Total number of micrographs | 32,886 (22,865 from monomer grids, 10,021 from dimer grids) | | |
| Selected micrographs | 26,686 (18,074 from monomer grids, 8,612 from dimer grids) | | |
| Initial particle images | 4,687,663 | | |
| Final particle images | 183,273 | | 55,178 |
| Starting model | *De novo* | | *De novo* |
| Map resolution | 3.92 | 3.78 | 6.38 |
| FSC threshold | 0.132 | 0.132 | 0.132 |
| Sharpening *B* factor ($Å^{-2}$) | 129.2 | 104.3 | 402.9 |
| EMDB code | EMD-28949 | EMD-28951 | EMD-28952 |
| **Model building and refinement** | | | |
| Initial models used | AF-Q38SD2-F1 | | |
| Map used for refinement | Composite map, EMD-28950 | | |
| Model resolution | 4.1 | | |
| FSC threshold | 0.5 | | |
| *Model composition* | | | |
| Non-hydrogen atoms | 11699 | | |
| Amino acid residues | 1472 | | |
| Protein molecules | 1 | | |
| *Real-space correlation* | | | |
| CCvolume | 0.72 | | |
| CCmask | 0.74 | | |
| Mean *B* factor ($Å^2$) | 94.14 | | |
| *RMS deviations* | | | |
| Bond lengths (Å) (outliers >4σ) | 0.004 (0) | | |
| Bond angles (°) (outliers >4σ) | 0.649 (1) | | |

| *Validation* | |
|---|---|
| *MOLProbity* score | 2.13 |
| Clashscore | 14.36 |
| Rotamer outliers (%) | 0.69 |
| CaBLAM outliers (%) | 4.63 |
| Cβ outliers (%) | 0.00 |
| *Ramachandran plot* | |
| Favored (%) | 92.42 |
| Allowed (%) | 7.51 |
| Outliers (%) | 0.00 |
| PDB code | 8FAC |

**Supplementary Movie Caption:** 3D variability analysis of LRRK1. Movie shows the three variability modes solved for LRRK1.

**Supplementary Material References:**

1 Myasnikov, A. *et al.* Structural analysis of the full-length human LRRK2. *Cell*, 1-9 (2021). https://doi.org:10.1016/j.cell.2021.05.004

2 Guo, Z., Hou, X., Goody, R. S. & Itzen, A. Intermediates in the guanine nucleotide exchange reaction of Rab8 protein catalyzed by guanine nucleotide exchange factors Rabin8 and GRAB. *Journal of Biological Chemistry* **288**, 32466-32474 (2013). https://doi.org:10.1074/jbc.M113.498329

3 Martinez Fiesco, J. A., Durrant, D. E., Morrison, D. K. & Zhang, P. Structural insights into the BRAF monomer-to-dimer transition mediated by RAS binding. *Nature Communications* **13**, 1-14 (2022). https://doi.org:10.1038/s41467-022-28084-3

4 Liau, N. P. D. *et al.* The molecular basis of JAK/STAT inhibition by SOCS1. *Nature Communications* **9**, 1-14 (2018). https://doi.org:10.1038/s41467-018-04013-1

5 Mao, C., Zhou, M. & Uckun, F. M. Crystal Structure of Bruton's Tyrosine Kinase Domain Suggests a Novel Pathway for Activation and Provides Insights into the Molecular Basis of X-linked Agammaglobulinemia. *Journal of Biological Chemistry* **276**, 41435-41443 (2001). https://doi.org:10.1074/jbc.M104828200

6 Knighton, D. R. *et al.* Crystal structure of the catalytic subunit of cyclic adenosine monophosphate-dependent protein kinase. *Science* **253**, 407-414 (1991). https://doi.org:10.1126/science.1862342

7 Xie, X. *et al.* Crystal structure of JNK3: A kinase implicated in neuronal apoptosis. *Structure* **6**, 983-991 (1998). https://doi.org:10.1016/S0969-2126(98)00100-2

8 Jeffrey, P. D. *et al.* Mechanism of CDK activation revealed by the structure of a cyclinA-CDK2 complex. *Nature* **376**, 313-320 (1995). https://doi.org:10.1038/376313a0

9 Nowakowski, J. *et al.* Structures of the cancer-related Aurora-A, FAK, and EphA2 protein kinases from nanovolume crystallography. *Structure* **10**, 1659-1667 (2002). https://doi.org:10.1016/S0969-2126(02)00907-3

10 Leonard, T. A., Róycki, B., Saidi, L. F., Hummer, G. & Hurley, J. H. Crystal structure and allosteric activation of protein kinase C βII. *Cell* **144**, 55-66 (2011). https://doi.org:10.1016/j.cell.2010.12.013

11 Deniston, C. K. *et al.* Structure of LRRK2 in Parkinson’s disease and model for microtubule interaction. *Nature* (2020). https://doi.org:10.1038/s41586-020-2673-2
